# IMPA1 dependent regulation of plasma membrane phosphatidylinositol 4,5-bisphosphate turnover and calcium signalling by lithium

**DOI:** 10.1101/2022.10.14.512101

**Authors:** Sankhanil Saha, Harini Krishnan, Raghu Padinjat

## Abstract

Lithium (Li) is a widely used as a mood stabilizer in the clinical management of Bipolar Affective Disorder (BPAD). However, the molecular targets of Li in neural cells that underpin its therapeutic effect remain unresolved. Inositol monophosphatase (IMPA1), is an enzyme involved in the resynthesis of phosphatidylinositol 4,5-bisphosphate (PIP_2_) following receptor-activated phospholipase C (PLC) signalling. *In vitro*, Li inhibits IMPA1, but the relevance of this inhibition within neural cells remains unknown. Here we report that in human cells, treatment with therapeutically relevant concentrations of Li reduces receptor activated calcium release from intracellular stores and delays the resynthesis of PIP_2_ following receptor activated PLC signalling. Both these effects of Li are abrogated in cells where IMPA1 has been deleted. We also observed that in human forebrain cortical neurons, treatment with Li results in reduced neuronal excitability as well as reduced calcium signals following receptor activated PLC signalling. Following Li treatment of human forebrain cortical neurons, transcriptome analyses reveal downregulation of multiple components of the glutamate receptor signalling system. Glutamate is a key excitatory neurotransmitter in the human brain and thus our findings provide an insight into the mechanisms underlying the dampening of neuronal excitability following Li treatment. Collectively, our findings suggest that Li inhibits receptor activated PLC signalling leading to an altered transcriptional response and reduced neuronal excitability.

## Introduction

Bipolar affective disorder (BPAD) is a human psychiatric illness characterized by disruptive mood swings; the defining characteristics being the alteration between mania and depression associated with an elevated rate of suicide among patients (Osby et al., 2001). BPAD has been regarded by the World Health Organization (WHO) as one of the leading causes of disability worldwide (Lopez and Murray, 1998). Mood stabilizers like lithium (Li), Carbamazepine and Valproate remain the first-line mode of medication for bipolar disorder. Among these drugs, Li stands out as the go-to drug for treating acute manic phases and for its preventive effect against relapses of manic episodes (Cipriani et al., 2013; Geddes et al., 2004). Although Li remains the mainstay in BPAD treatment, individual patient responses towards Li are variable-it remains ineffective for a large proportion (approximately 30%) of BPAD patients, the so-called ‘Li non-responders’, whereas another 30% of patients are only partially responsive to Li (Hou et al., 2016). The reason for this variable response is unknown and remains a topic of intense interest. Despite substantial research over several decades, the molecular and biochemical mechanism by which Li exerts its effects on human brain cells that are relevant to functional improvements remains unclear (Haggarty et al., 2021). Many molecular targets for Li have been described (Roux and Dosseto, 2017). These include Inositol monophosphatase (IMPase) (Harwood, 2005), Glycogen synthasekinase-3β (GSK-3β) (Freland and Beaulieu, 2012), Sodium myo-inositol co-transporter (SMIT1) (Dai et al., 2016) and Bisphosphate-3’-nucleotidase (BPNT1) (Spiegelberg et al., 2005). The two most well studied targets of Li are Inositol monophosphatase (IMPase) and Glycogen synthase kinase-3β (GSK-3β).

During GPCR signalling, many receptors activate phospholipase C (PLC) leading to the hydrolysis of the signalling lipid phosphatidylinositol 4,5-bisphosphate (PIP_2_) generating inositol 1,4,5-trisphosphate (IP_3_) and diacylglycerol (DAG) (Berridge, 2009), ultimately leading to changes in intracellular calcium [Ca^2+^]_i_. This mechanism of cell signalling is used by numerous G-protein coupled receptors in the brain that participate in synaptic transmission such as glutamate and acetyl choline. Previous studies in rat hippocampal cultures has shown that treatment with Li diminishes Ca signalling responses to Gq-PLC coupled neurotransmitter receptors such as the metabotropic glutamate receptor and muscarinic acetylcholine receptor (Sourial-Bassillious et al., 2009)and alterations in [Ca^2+^]_i_ have been reported in cell lines derived from BPAD patients (Wasserman et al., 2004). Most recently, a meta-analysis has revealed that basal levels of [Ca^2+^]_i_ as well as [Ca^2+^]_i_ responses to agonist stimulation in BPAD patients are elevated (Harrison et al., 2021). Thus, a link between Ca signalling and BPAD has been previously noted although the underlying mechanism in this context remains unclear.

A long-standing hypothesis for the therapeutic effect of Li is its impact on inositol lipid turnover during receptor activated G-protein coupled signalling through inhibition of IMPase (Harwood, 2005). PIP_2_ is a low abundance membrane lipid and its resynthesis is required to sustain Ca signalling during high rates of PLC activity as is seen in the brain (Yang et al., 2016) during the activation of many GPCR required for neurotransmission such as the metabotropic glutamate receptor (Reiner and Levitz, 2018). The resynthesis of PIP_2_ occurs via a set of biochemical reactions referred to as the PIP_2_ cycle (Cockcroft and Raghu, 2016). The sequential dephosphorylation of IP_3_ produced during PLC activity to generate inositol, is required for PIP_2_ resynthesis (Fig 1A). IMPase is the enzyme required for the final step in the conversion of inositol 1-phosphate to inositol. Biochemical analyses showed a decrease in myo-inositol concentration in cerebral cortex of lithium-treated rats accompanied by an increase in the concentration of inositol monophosphate suggesting that Li might inhibit IMPase (Allison et al., 1976; Allison, J.H., 1971). Subsequent studies showed that Li binds to IMPase and inhibits the enzyme in a non-competitive manner (Allison et al., 1976). These observations led Berridge to propose the “Inositol depletion hypothesis” (Berridge et al., 1989) which posits that during PLC signalling in the brain, the inhibition of IMPase by Li restricts the supply of inositol for PIP_2_ resynthesis leading to down regulation of neurotransmitter receptor signalling, thus reducing excitability in the brain and the control of mania. However, the link between G-protein coupled PIP_2_ turnover, [Ca^2+^]_i_ signalling and the action of Li in human neurons remains to be established.

**Figure 1:**
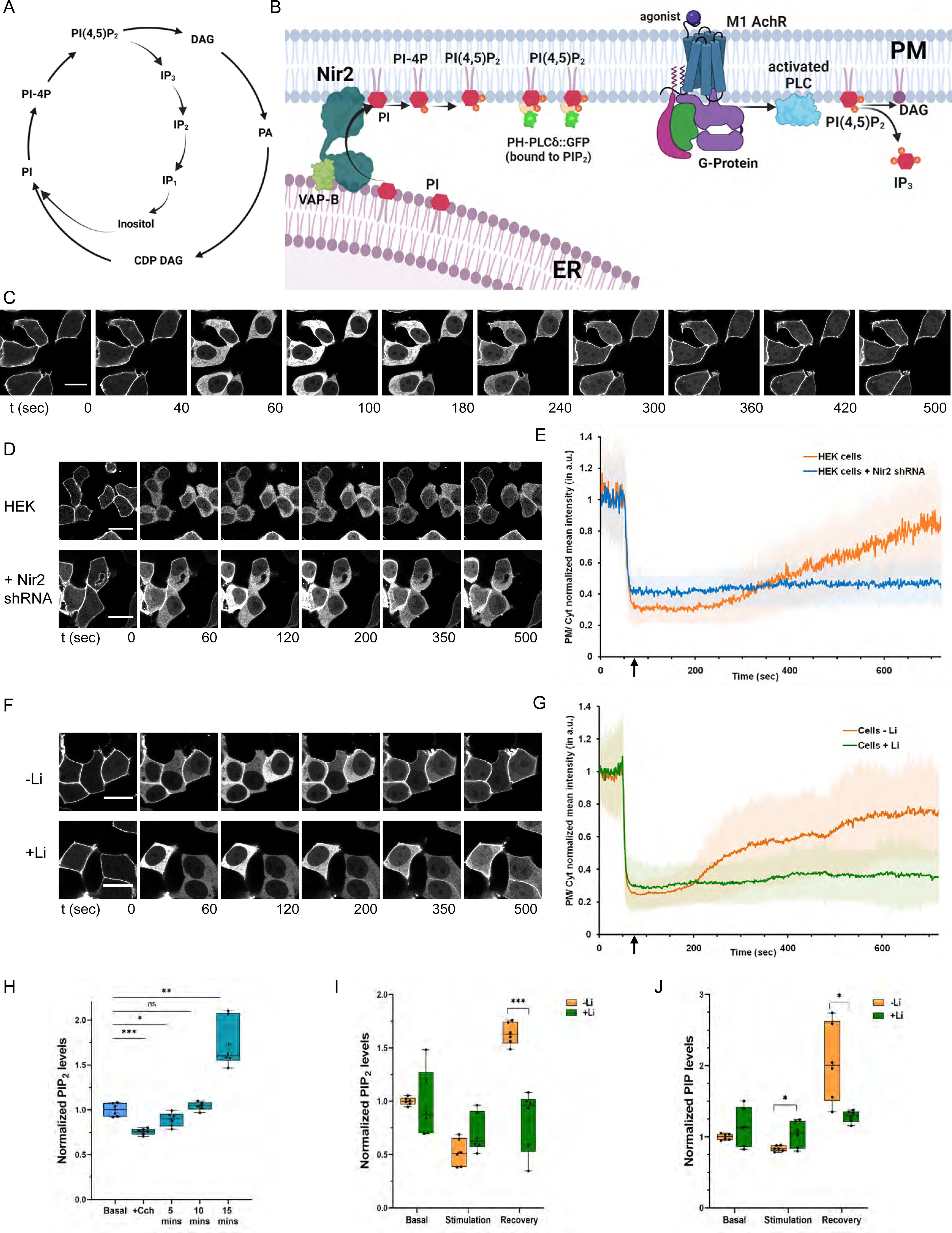
An *in vivo* model system to study the impact of Li on PLC induced PIP_2_ turnover. (A) The Phosphatidylinositol bisphosphate (PIP_2_) cycle. (B) Schematic representation of an *in vivo* system elucidating the role of Nir2 and m1AchR in the PIP_2_ cycle and the PH-PLCδ::GFP probe binding to the PIP_2_ at the plasma membrane. (C) The PH-PLCδ::GFP probe translocates to the cytoplasm and back to the plasma membrane, corresponding to the hydrolysis and regeneration of PIP_2_ respectively (control-M1; HEK293T cells expressing m1AchR). (D, E) Nir2 depletion leads to a delayed regeneration of PIP_2_ at the plasma membrane. Changes in membrane localization of the PH-PLCδ::GFP probe after Cch stimulation (denoted by the ↑), are quantified via the fluorescence intensity ratio of the plasma membrane and the cytosol (PM/ Cyt) at various times. This ratio is normalized to the first time-point and the mean ± SD is plotted (from three experiments each performed in replicates) for control cells (orange line, n =33), and for cells treated with Nir2 shRNA (blue line, n =28). (F, G) Rate of regeneration of PIP_2_ at the plasma membrane is shown for control-M1 cells (HEK293T cells expressing m1AchR) subjected to lithium treatment, compared to untreated cells. Mean ± SD is plotted from four experiments, each performed in replicates (untreated cells-orange line, n = 58; cells treated with Li-green line, n =53). [Scale Bar-20 μM]. (H) Total PIP_2_ levels recovered after various time intervals post stimulation with Cch. (I, J) Total PIP_2_ and PIP levels using LCMS from whole cell lipid extract of untreated and lithium treated cells (n=6 for both). [Statistical test: (H) one-way ANOVA with post hoc Tukey’s multiple pairwise comparisons. *p value < 0.05; **p value <0.01; ***p value < 0.001; ****p value < 0.0001 and (I, J) Multiple unpaired t test. *p value < 0.05; **p value <0.01; ***p value < 0.001].

The mammalian genome includes two genes encoding IMPase activity, IMPA1 and IMPA2. However, IMPA1 is more abundant in the human brain (Schubert et al., 2015)and is inhibited by Li at therapeutically relevant concentrations (Ohnishi et al., 2007). *In vivo* studies have also suggested a role of Li in regulating IMPase function. Briefly, (i) in rodent models, Li treatment reduced brain inositol levels and elevated Inositol monophosphate (IP_1_), the substrate of IMPase in the cerebral cortex of rat (Sherman et al., 1981) (ii) treatment of human BPAD patients with Li led to a reduction in PIP_2_ levels in platelet membranes (Soares et al., 1997; Soares et al., 1999; Soares et al., 2000). However, there are also findings that do not support this hypotheses (i) IMPA1 knockout mice showed only a modest reduction in brain inositol levels (Cryns et al., 2008) and the impact of IMPA1 knockout on receptor activated PIP_2_ turnover remains unknown, (ii) genome-wide association studies have not revealed polymorphisms in IMPA1 that are linked to BPAD or Li sensitivity (Hou et al., 2016; Sjoholt et al., 2000).

In this study, we have tested the hypothesis that treatment of human cells with Li diminishes calcium signalling. We find that this change is underpinned by reductions in G-protein coupled PIP_2_ turnover. These changes in calcium signalling were also observed in human iPSC derived forebrain cortical neurons and accompanied by reduction in neuronal excitability. Transcriptomic analysis revealed that Li treatment exerts its effect on neuronal excitability through multiple mechanisms including but not restricted to GPCR activated calcium signalling. Our findings open new approaches to predict Li responsiveness in BPAD patients and develop novel strategies for the clinical management of this disorder.

## Materials and Methods

### DNA constructs

For PIP_2_ measurement, PH-PLCδ::GFP probe was used which was a gift from Tamas Balla (Addgene plasmid # 51407). The cDNA for human muscarinic acetylcholine receptor m1AchR was obtained from cDNA Resource Center (MAR0100000) and was cloned via Gibson assembly into a lentiviral pHR transfer backbone (having Puromycin resistance cassette) for lentiviral generation. The Nir2 shRNA was obtained from GE Dharmacon (RHS4430-200186441: Clone id V2LHS_62836).

### Mammalian cell culture, treatment and transfection

HEK293T cells were maintained under standard conditions at (37°C, with 5% CO_2_) in DMEM high Glucose media (Dulbecco’s Modified Eagle Medium; Life Technologies), supplemented with 10% Foetal bovine serum. For Li treatment, the cells were grown and maintained in this media, supplemented with 1 mM LiCl (Sigma) for 5 days (unless mentioned otherwise). The cells were tested for mycoplasma, prior to the experiments. MycoAlert™ (Lonza) was used per the manufacturer’s protocol to test the spent media for mycoplasma contamination. Transfection with the plasmid expressing PH-PLCδ::GFP probe was done 24 hours before the imaging experiments using Polyethylenimine solution (PEI; Polysciences Inc.; 1 mg/ml) when the cells were around 70% confluent.

### Lentiviral transduction to generate HEK293T cell line stably expressing muscarinic acetylcholine receptor m1AchR

Recombinant Lentiviral particles were packaged in HEK293T cells grown in DMEM high Glucose media, supplemented with 10% FBS. The HEK293T cells cultured in the 6 well plates were transfected with a solution consisting of 1.2 μg of pHR-IRES plasmid encoding muscarinic acetylcholine receptor *CHRM1* cDNA, along with 1.2 μg of each of the helper plasmids (psPAX and pMD2.G) in 500 μl of Opti-MEM (Invitrogen) and 110 μl Polyethylenimine (PEI, 1 mg/ ml). Medium was changed after 12 hours and the virus was harvested at 72 h after transfection. Then HEK293T cells were transduced with these recombinant viral particles and selected via Puromycin treatment (1.5 μg/ ml) for 1 week-to obtain the HEK293T cell line, expressing m1AchR. Western Blot confirming the expression of m1AchR is shown in supplementary figures (Supp. Fig 3C, D).

### Generation of Neural Stem Cells and differentiation and maintenance of neuronal culture

Neural Stem Cells (NSCs) were generated as previously described (Mukherjee et al., 2019), with slight modifications. Human iPSCs were differentiated to embryoid bodies in E6 medium (Gibco). After that, primary neural rosettes that were formed, were manually picked, passaged, and eventually plated as NSC monolayer in Neural Expansion Medium (NEM; DMEM/F12 with 2mM Glutamax, supplemented with N2, B27 without Vitamin A, 20 ng/ml bFGF, 100 μM non-essential amino acids, 2 µg/ml Heparin Sulphate, 100 U/ml penicillin-streptomycin) (all components procured from Life Technologies). The NSCs were subjected to CD133+ selection as previously described via fluorescence-activated cell sorting (FACS) (Peh et al., 2009) to eliminate cells of other lineages. The sorted NSCs were further characterized by immunofluorescence for the specific markers. Neural Stem Cells (NSCs) were grown on Matrigel coated tissue dishes in NEM at 37°C and 5% CO_2_.

For differentiation, NSCs were seeded onto Matrigel-Growth Factor Reduced (Corning) coated cover-slip dishes or 35 mm dishes and expanded in NEM till the cells reached 70% confluency. NEM was then withdrawn and replaced with Neural Differentiation Medium (NDM; DMEM/F12 with 2mM Glutamax, supplemented with N2, B27 containing Vitamin A, 100 μM non-essential amino acids, 100 U/ml penicillin-streptomycin, 10 ng/ml BDNF, 10 ng/ml GDNF, 10 ng/ml IGF-1, 50 μg/ml Ascorbic acid (Sigma) and 2 µM DAPT (Sigma)). DAPT was supplemented for the first 14 days. Media was changed every 2 days during the differentiation process. All the experiments were done 45 DIV post differentiation (The day of addition of NDM was DIV 0). For chronic Li treatment, the neuronal cultures were maintained in NDM supplemented with 1 mM LiCl for the last 10 days (DIV35 to DIV45).

### Live cell imaging for the PIP_2_ probe

To analyse the dynamics of PIP_2_ at the plasma membrane by following the change in translocation of the PH-PLCδ::GFP probe, live cell imaging was performed as stated in *Varnai et al* (Várnai and Balla, 1998), with minor modifications. 20000 cells were seeded in Poly-L Ornithine (PLO) coated cover-slip dishes and grown for 96 hours with or without Li. 24 hours prior to imaging, the cells were transfected with PH-PLCδ::GFP plasmid using Polyethylenimine as per manufacturer’s protocols. Carbamoylcholine chloride mediated changes in the membrane localization of PH-PLCδ::GFP probe was examined by a time-lapse method using an Olympus FV 3000 confocal microscope. The cells were kept in pre-warmed Kreb’s Ringer Buffer (120 mM NaCl, 4.7mM KCl, 1.2 mM CaCl_2_, 0.7 mM MgSO_4_, 10 mM Glucose, 10 mM HEPES; pH-7.4) and at 37°C while imaging. Changes in the PIP_2_ membrane localization were quantified by taking a fluorescence intensity ratio of the plasma membrane (F_PM_) and the cytosol (F_Cyt_) across the time-lapse and normalizing it to the first value.

### Calcium Imaging for Carbamoylcholine stimulation

For studying the calcium physiology, HEK293T-M1 cells were seeded in Poly-L Ornithine (PLO) coated cover-slip dishes while iPSC derived cortical neurons (DIV45) were grown on Matrigel coated cover-slip dishes. Before imaging, the cells were washed twice with Hank’s Buffered Salt Solution (HBSS) (10 mM HEPES, 118 mM NaCl, 4.96 mM KCl, 1.18 mM MgSO_4_, 1.18 mM KH_2_PO_4_, 10 mM Glucose, 2 mM CaCl_2_; pH-7.4) and then loaded with the ratiometric Ca^2+^ indicator Fura-2 AM (4 µM) (Acetoxymethylester, Invitrogen) along with 0.002% Pluronic F-127 for 30 minutes. The excess dye was removed by washing the cells thrice with HBSS and cells were incubated for an additional 20 minutes to equilibrate the intracellular dye concentration and allow de-esterification. Imaging was performed for 10 min using a 40X objective (N.A. 0.75) of wide-field fluorescence microscope Olympus IX-83 for HEK293T cells, while a 20X objective (N.A. 0.50) was used for neuronal cultures. Free- and Ca^2+^-bound Fura-2 fluorescence intensities (excitation 340/380 nm, emission 510 nm) were recorded at every 5 seconds intervals using an EM-CCD camera (Evolve 512 Delta, Photometrics). Basal cytosolic Ca^2+^ was recorded in HBSS buffer for the initial 18 frames. Carbamoylcholine chloride (20 µM and 50 µM respectively for HEK and neuronal cultures, as determined in preliminary concentration– response experiments) (Sigma Aldrich-C4382) was added to the cells for activating PLC and thereby, inducing Ca^2+^ release from the endoplasmic reticulum store. After recording for 42 frames, 20 μM Ionomycin (Calbiochem) was added and recorded for 24 frames for recording R_max_. 24 more frames were recorded, post addition of 4 mM EGTA (containing 0.01% Triton-X-100) for chelation of Ca^2+^ ions and determination of R_min_.

### Calcium Imaging for SOCE (Store Operated Calcium Entry)

For the SOCE experiment, the cells were kept in ‘zero Ca^2+^ HBSS’ (10 mM HEPES, 118 mM NaCl, 4.96 mM KCl, 1.18 mM MgSO_4_, 1.18 mM KH_2_PO_4_, 10 mM Glucose; pH-7.4) just before the imaging. After recording the initial 12 frames for basal cytosolic Ca^2+^, an optimal concentration of Thapsigargin (Tg) (10 μM) (Invitrogen) was added for the store depletion and 60 frames were recorded. Then, 2 mM CaCl_2_ was then added to the extracellular buffer to induce SOCE-60 frames were recorded post induction of SOCE. Subsequently, 20 μM Ionomycin (Calbiochem) was added and recorded for 24 frames; then 4 mM EGTA (containing 0.01% Triton-X-100) was added and 24 frames were recorded. For studying PLC activated calcium mobilization from stores and SOCE, Carbamoylcholine chloride (20 µM and 50 µM respectively for HEK and neuronal cultures) was added instead of Thapsigargin.

### Analysis of Calcium recordings

The CellSens Olympus software was used for the analysis. A dynamic region of interest (ROI) was drawn for each cell which exhibited rise in [Ca^2+^]_i_ (as seen by increase in fluorescence post-stimulation) to track the fluorescence changes over time. The emission intensities corresponding to 340 nm and 380 nm excitations were used to calculate the 340/380 ratio for each cell across all time points. The [Ca^2+^]_i_ was estimated using the Grynkiewicz equation (Grynkiewicz et al., 1985) as follows:

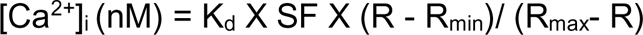

where R_min_ and R_max_ refer to the minimum and maximum 340/380 ratio respectively. 224 nM was taken as K_d_ for Fura-2 AM in human cells and SF (scaling factor) was calculated for each cell by dividing the fluorescence emission intensity at Ca^2+^ free form with the fluorescence emission intensity at Ca^2+^ bound form of the dye after excitation at 380 nm. The final Ca^2+^ traces were plotted using GraphPad Prism 9.0.

### Generation of the IMPA1 deletion in cell lines

For generation of HEK293T-IMPA1^-/-^ cell line, the initial region of the coding sequence (CDS) (exon1) of the IMPA1 gene was targeted by a pair of guide RNAs (sgRNAs) and deleted via CRISPR-Cas9 technology, then these cells were transduced to stably express Muscarinic receptor M1. The sgRNAs targeting the start of the CDS of IMPA1 gene were designed via the <u>crispr.mit.edu</u>. The sequences of the sgRNAs are as follows:

5’ GGAGAACCACCTTGTTGGCCAAGCTGGAATGTAGTGGCGTTTTAGAGCTAGAAATAGCAAGTT 3’

5’ GGAGAACCACCTTGTTGGGTAATATGGTACAGACACACGTTTTAGAGCTAGAAATAGCAAGTT 3’

These sgRNAs were cloned individually into humanized pgRNA plasmids [pgRNA-humanized was a gift from Stanley Qi (Addgene plasmid # 44248)]. The cells were co-transfected with the three plasmids (both the pgRNA humanized plasmids encoding the sgRNAs and pLentiCas9 blast encoding for the Cas9 protein endonuclease from *Streptococcus pyogenes*) and selected with Puromycin (2 μg/ ml) for 48 hours. Then the genomic DNA was isolated, and a Surveyor PCR was performed to check the occurrence of the deletion. Once confirmed that the sgRNAs were able to target the exon 1 of IMPA1 gene, the transfected cells were seeded in 96 wells as single cells and gradually expanded. After selection via Surveyor PCR using Surveyor primers (primers flanking the region ought to be deleted), a HEK293T cell line was selected which had showed only the shorter amplicon (amplicon after deletion) in the Surveyor PCR. This HEK293T cell line was then expanded and the expression of IMPA1 was checked by Western blotting using the anti-IMPA1 antibody. No corresponding bands for IMPA1 were observed, indicating that IMPA1 was knocked out in this cell line. This HEK293T-IMPA1^-/-^ cell line did not show any morphological difference with respect to HEK293T cells and did not require inositol supplementation in the media for maintenance.

### Western Blotting

For estimating the expression of the IMPA1, cells were suspended in appropriate volume of lysis buffer and kept on ice for lysis; 10 μl of it was used for Bradford protein assay (BioRad) for quantification. Post quantification, equal amount of samples was heated at 95°C with Laemmli loading buffer for 5 min, then loaded and separated on 10% SDS-PAGE gel. For estimating the expression of the muscarinic acetylcholine receptor m1AchR, cells were pelleted and directly lysed by adding Laemmli’s loading buffer and repeated syringing with insulin syringes (31 gauge). Then the samples were heated at 95°C for 5 min and loaded onto a pre-cast Polyacrylamide 4-12% gradient gel (Bolt, Invitrogen). The proteins were then transferred onto a nitrocellulose membrane and then blocked with 10% Blotto (Santa Cruz Biotechnology) diluted in Tris Buffer Saline containing 0.1% Tween-20 (0.1% TBST) for 40 minutes. Subsequently the membrane was incubated with the rabbit anti-IMPA1 antibody (1:10000 diluted in 5% BSA in 0.1% TBST) (Abcam ab184165) or goat anti-CHRM1 antibody (1 µg/ ml, diluted in 5% BSA in 0.1% TBST) (Abcam ab77098) overnight at 4°C. For loading control, the blot was incubated with mouse anti-β-Tubulin antibody (1:4000 diluted in 5% BSA in 0.1% TBST) (DSHB Hybridoma Product E7). The blots were then washed thrice with 0.1% TBST and then incubated with the appropriate HRP-conjugated secondary antibodies (1:10000 dilutions; Jackson Laboratories, Inc.) for 1.5 hrs. Post incubation with the secondary antibodies, the blot was washed thrice with 0.1% TBST and developed using Clarity Western ECL substrate (Biorad) on a GEImageQuant LAS 4000 system. The ImageJ software was used for the densitometric analysis of the blots. Firstly, the background intensities were subtracted from the images using an average of mean background intensities of a few ROIs adjacent to the band of interest. Then ROIs were drawn around the bands of interest and the integrated density of each ROI was extracted from the image. Finally, the ratio of the integrated density of the IMPase to that of the loading control (β-Tubulin) was calculated.

### Calcium imaging for measuring transients

For studying the effect of Li on the excitability of cortical neurons, spontaneous calcium events (calcium transients) were measured in D149 cortical neuronal cultures at 45 DIV for both untreated and Li treated neurons, according to previously published lab protocol (Sharma et al., 2020). The neuronal cultures were washed twice for 10 minutes with HBSS and then loaded with 4 μM of Fluo-4 AM (Molecular probes, F14201) in HBSS having 0.002% Pluronic F-127 for 30 min at room temperature. Imaging was performed for a time-span of 10 min using a 20X objective (N.A. 0.50) with a time interval of 1 second at 488 nm illumination. The baseline measurement was recorded for a period of 4 min to visualize the calcium transients. Then 10 µM Tetrodotoxin (TTX, Hellobio) was added to abolish calcium transients and recorded for another 4 min. A dynamic region of interest (ROI) was manually drawn around each neuronal soma; individual Ca^2+^ traces were then obtained using the CellSens Dimensions software. The fluorescence intensity value from each neuronal soma was normalized to the first baseline signal and the individual traces were plotted using GraphPad Prism 9.0. The frequency of calcium transients was measured manually from first-derivative filter traces.

### Post-hoc immunofluorescence for neuronal characterization

Post-calcium imaging, the neuronal cultures were washed thoroughly a few times with HBSS buffer and then fixed with 4% PFA dissolved in PBS for 20 mins. Post-fixation, the cultures were washed twice with PBS for 5 min each and then permeabilized with 0.1% Triton-X (dissolved in PBS). The cultures were then blocked with 5% BSA diluted in PBS, for 1 hr at room temperature. Subsequently the neurons were incubated with the Primary antibodies diluted in 5% BSA in PBS overnight at 4°C. Next day, the cultures were subjected to three washes with PBS (10 minutes each) to remove non-specifically bound primary antibody. Respective secondary antibodies at a dilution of 1:300 in 5% BSA in PBS were used for incubation for 2 hours. DAPI was used as a nuclear marker at a dilution of 1:1000.

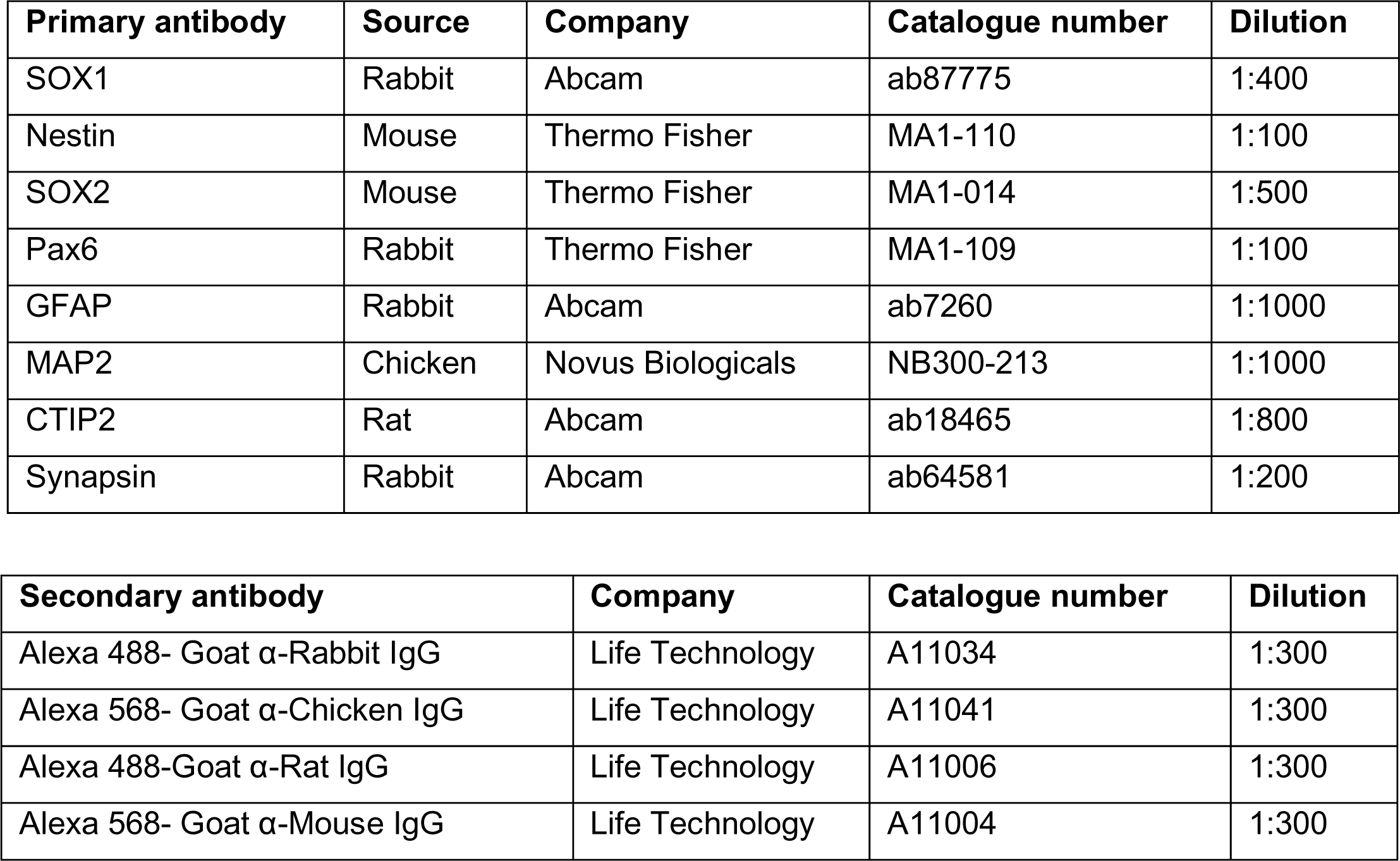

### RNA isolation, cDNA synthesis and Real-Time Quantitative PCR

The total RNA was extracted from HEK293T cells or mature neuronal cultures, in multiple biological replicates using TRIzol (Ambion, Life Technologies) following manufacturer’s protocol and thereby, quantified via Nanodrop 1000 spectrophotometer (Thermo Fisher Scientific). For cDNA synthesis, 1 μg of the RNA from each replicate was treated with DNase I (amplification grade, Thermo Fisher Scientific) and then incubated with Superscript II reverse transcriptase (Invitrogen) along with random hexamers and dNTPs. A no reverse transcription control sample was also included for each sample. Real-Time Quantitative PCR analysis was performed on an Applied Biosystems 7500 fast qPCR system using diluted cDNA samples and primers against genes of interest and GAPDH as internal controls. The C_t_ values obtained for different genes were normalized to those of GAPDH from the same sample. The relative expression levels were calculated using ΔC_t_ method, whereas the fold change was calculated using ΔΔC_t_ method. The primers used for qRT-PCR are listed in the Supplementary Table 2.

### LCMS based PIP and PIP_2_ measurements

To validate whether the PH-PLCδ-GFP probe based imaging of PIP_2_ in the HEK293T cells reflected the changes in PIP_2_ level, mass spectrometry measurements of PIP_2_ from the cells was carried out following existing lab protocols (Sharma et al., 2019). Reverse phase liquid chromatography was coupled with high sensitivity mass spectrometry (LCMS) and a Multiple Reaction Monitoring (MRM) method was employed to detect PIP_2_ levels from the whole cells.

#### Lipid extraction

Equal number of cells from a single well of a 12 well plate (counted by a haemocytometer while seeding) were pelleted and then gently resuspended into 170 μl of 1X PBS in a 2 ml low-bind polypropylene centrifuge tube. To this, 750 µl of ice-cold quench mixture (MeOH/CHCl_3_/1M HCl in the volumetric ratio of 484/242/23.55), followed by 15 µl of a pre-mixed ISD (Internal Standard) mixture containing 25 ng of 37:4 PIP [**PI 4P 17:0/20:4 (LM-1901)]**, 25 ng of 37:4 PIP_2_ [**PI 4,5P_2_ 17:0/20:4 (LM-1904)]** and 50 ng of 31:1 Phosphatidylethanolamine [**PE 17:0/14:1 (LM-1104)]** was added (All the lipid standards were procured from Avanti Lipids). For Carbamoylcholine chloride mediated PLC activation, cells were suspended in 85 μl of 1XPBS and then 85 μl of 20 μM Carbamoylcholine chloride (made in PBS) (final concentration – 10 μM) was added for 1 min at 37°C in shaking conditions. For the regeneration of PIP_2_ post PLC mediated hydrolysis, 1 ml of ice-cold PBS was added to dilute the agonist Carbamoylcholine chloride and stop the stimulation. Cells were pelleted and washed and then resuspended in pre-warmed buffer (37°C) and kept for 15 mins for recovery (time given for regeneration of PIP_2_). Post 15 mins of incubation, cells were pelleted and then resuspended in 170 μl of PBS, to which 750 μl of quench mixture and 15 µl of a pre-mixed ISD were added as stated previously. The mixture was vortexed and 725 μl of CHCl_3_ and 170 µl of 2.4 M HCl were added. After vortexing again for 2 min at 1500 rpm, the phases were separated by centrifugation for 3 min at 1500 g. The lower organic phase was then collected and added to fresh tubes containing 708 µl of Lower Phase Wash Solution (LPWS; MeOH/1M HCl/CHCl_3_ in the volumetric ratio of 235/245/15). The tubes were then vortexed and spun at 1500 rpm again for 3 min to separate the phases. The resultant lower organic phase was collected into a fresh tube and subjected to the Derivatization reaction.

#### Derivatization of Lipids

50 μl of 2M TMS-Diazomethane was added to the collected organic phase (TMS-Diazomethane is toxic and is to be used with utmost precautions and following safety guidelines). The reaction was allowed for 10 min at 600 rpm at room temperature, post which 10 μl of Glacial acetic acid was added to quench the reaction. 500 μl of Post Derivatization Wash Solution (PDWS-CHCl_3_/MeOH/H_2_O in the volumetric ratio of 24/12/9; the upper phase was used) was then added to the sample, vortexed and spun down at 1500 g for 3 min. The upper aqueous phase was discarded, and the wash step was repeated. To the final organic phase, 45 µl MeOH and 5 µl H_2_O was added, mixed, and spun down. The samples were then dried in a SpeedVac at 400 rpm for 150 mins under vacuum and 90 µl of MeOH was added to reconstitute the sample to a final volume of about 100µl.

#### Chromatography and Mass spectrometry

The chromatographic separation was performed on an Acquity UPLC BEH300 C4 column (100 × 1.0 mm; 1.7 µm particle size - Waters Corporation, USA) using a Waters Aquity UPLC system connected to an ABSCIEX 6500 QTRAP mass spectrometer for ion detection. All the samples were injected in duplicates and the flow rate was set to 100 µl /min. Solvent gradients were set, starting from 55% Solvent A (Water + 0.1% Formic Acid)-45% Solvent B (Acetonitrile + 0.1% Formic acid) from 0-5 min; then 45% B to 100% B from 5-10 min; 100% B was held from 10-15 min; between 15-16 min, 100% B was lowered to 45% B and held there till 20th min to re-equilibrate the column. On the mass spectrometer, Neutral Loss Scans was employed during pilot standardization experiments on biological samples to identify the parent ions that would lose neutral fragments corresponding to 490 a.m.u and 382 a.m.u-these were indicative of PIP_2_ and PIP species respectively and likewise 155 a.m.u for PE species (Sharma et al., 2019). Thereafter, we quantified PIP, PIP_2_ and PE species in biological samples using the selective Multiple Reaction Monitoring (MRM) method in the positive ion mode. Area under the peaks was calculated via the Sciex MultiQuant software. For each (run, area under the peak for each species of PIP_2_, PIP and PE was normalized to PIP_2_, PIP and PE internal standard peak respectively. Thereafter, the sum of normalized areas for all the species of PIP_2_ or PIP was taken and divided by the sum of normalized areas for all the species of PE in each of the biological samples to account for differences in total phospholipid extracted across samples. The MRM mass pairs used for PIP, PIP_2_ and PE species identification and quantification are listed below:

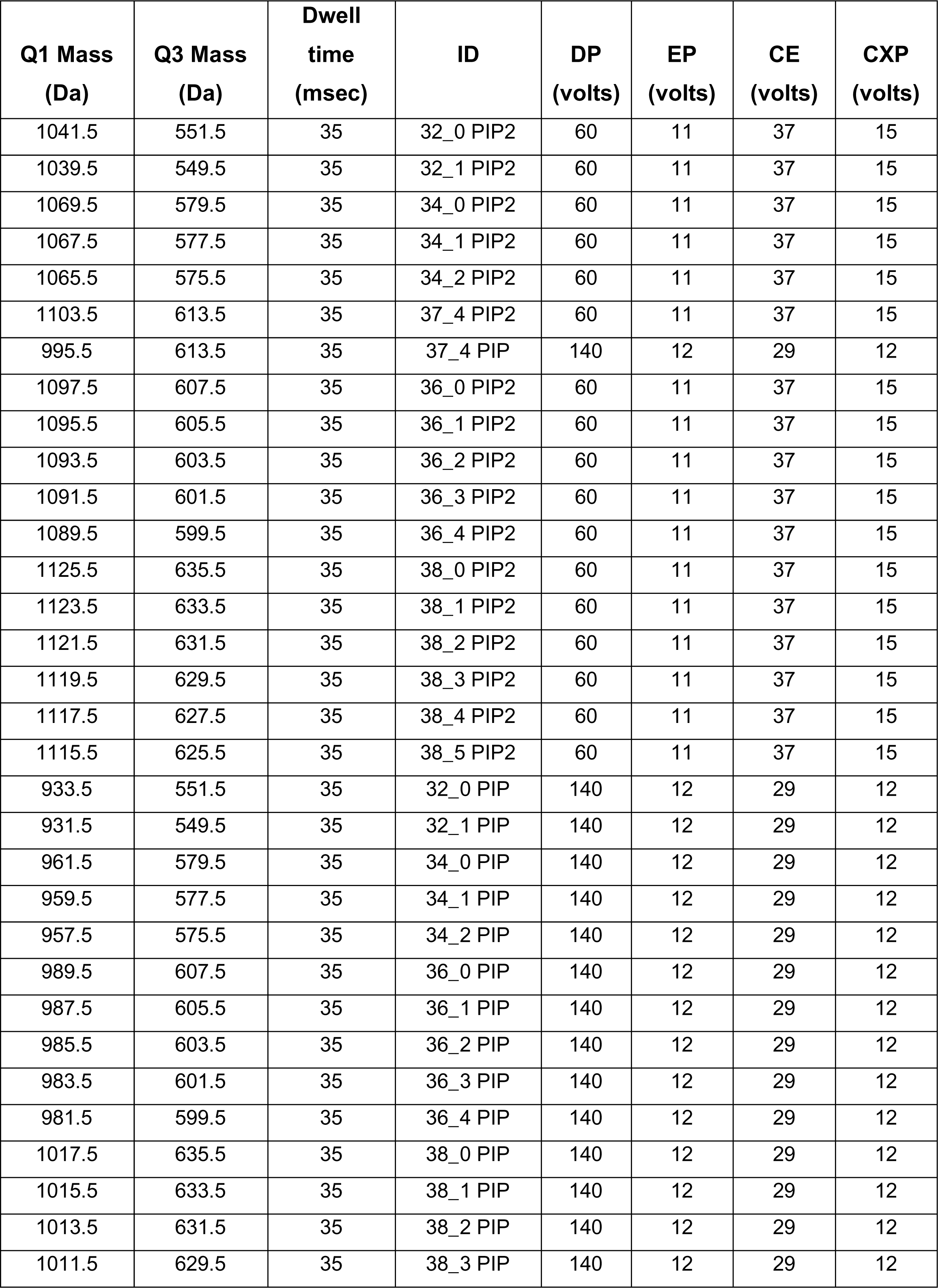
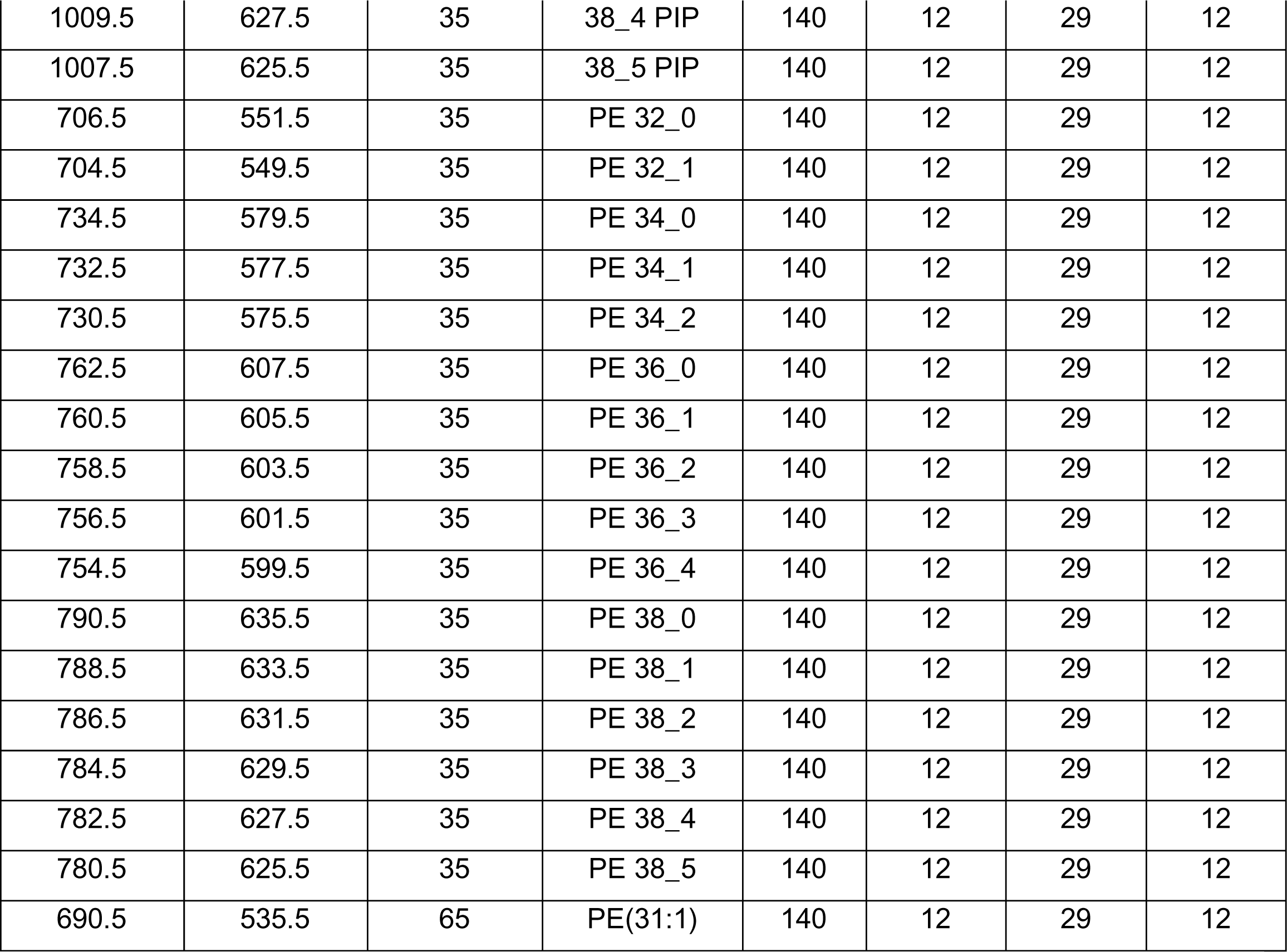

The other mass spectrometer parameters were:

ESI voltage: +5100-5200 V; Source Temperature: 300 C, Curtain Gas (CUR): 35-37, Ion Source Gas 1 (GS1):15-20, Ion Source Gas 2 (GS2): 15-20.

### RNA sequencing and transcriptomic analysis

RNA extracted from mature neurons (45DIV with and without Li treatment) and mature neurons (DIV45 with and without CHIR99021 treatment) as mentioned above was submitted for paired-end sequencing. Quality check and sequencing was carried out at the NCBS Sanger sequencing facility using Illumina Hiseq 2500 system. Single stranded RNA sequencing libraries were prepared for sequencing using NEBNext® Ultra™ II Directional RNA Library Prep with Sample Purification Beads (Catalog no-E7765L). Paired-end reads, (125 base pair for each neuronal sample and 100 base pair for each NSC samples) was obtained after sequencing. Post adapter trimming, FastQC was used to check the quality of reads. The reads were then aligned to human genome (GRCh38) using hisat2-build and hisat2-align module. The module for single strand sequencing was used in the Hisat-2 protocol. The output generated in the bam format, was further used to obtain the counts of each of the individual gene using HT-Seq pipeline. The differential expression of genes was then calculated using DeSeq2. GO annotation analysis was done using the DAVID tool.

### Sampling and statistical analysis

Each experiment was performed unblinded on different biological groups with multiple biological replicates. No statistical analysis was done a priori to determine the sample sizes. For the calcium imaging experiments, quantification was from a minimum of 100 cells for all genotypes from multiple dishes of each type. For the PIP_2_ probe experiments, approximately 50 cells were taken for the quantitative analysis. Statistical significance was computed using Student’s unpaired t-test to check for differences in means between samples of different genotypes. Two-way ANOVA with post hoc Tukey’s pairwise comparison was used for the grouped analysis. All statistical analyses were performed on Graph Pad Prism (version 9) and schematic representations were created with biorender.com.

## Results

### An *in vivo* model system to study the impact of Li on PLC induced PIP_2_ turnover

To study the turnover of PIP_2_ following receptor activated Phospholipase C (PLC) turnover, we created a cell line which heterologously expressed two components. To stimulate high levels of PLC activity, we expressed the human Muscarinic acetylcholine receptor (m1AchR) encoded by the gene *CHRM1*; previous studies have shown that activation of heterologously expressed m1AchR results in high rates of PIP_2_ hydrolysis leading to IP_3_ production (Parker et al., 1991; Raghu et al., 1997). To monitor the levels of PIP_2_ at the plasma membrane we expressed the PH-domain of PLC-δ fused to green fluorescent protein (PH-PLCδ::GFP). This probe binds to PIP_2_ and in imaging experiments; its relative distribution between the plasma membrane and the cytosol has been used by us and others to monitor plasma membrane PIP_2_ levels (Chakrabarti et al., 2015; Várnai and Balla, 1998). A cell line expressing both m1AchR and PH-PLCδ::GFP can therefore be used to stimulate PLC and monitor the turnover of PIP_2_ at the plasma membrane. We generated stable HEK293T cell lines expressing m1AchR (control-M1) and PH-PLCδ::GFP was transiently expressed in those cells (Fig 1B); when stimulated with the m1AchR agonist Carbamoylcholine (carbachol: Cch), the levels of PH-PLCδ::GFP at the plasma membrane drop following which they gradually recover back towards pre-stimulation levels (Fig 1C). The changes in plasma membrane PIP_2_ was quantified as the ratio of fluorescence intensity of the PH-PLCδ::GFP probe at the plasma membrane to its fluorescence intensity in the underlying cytosol in the same part of the cell. As a positive control for our assay to monitor PIP_2_ regeneration following PLC activation, we used shRNA to deplete Nir2; Nir2 (PITPNM1/RdgBαI) encodes for a phosphatidylinositol transfer protein required for the resynthesis of PIP_2_ following PLC stimulation (Chang and Liou, 2015; Kim et al., 2013; Kim et al., 2016). Under our experimental conditions, we found that depletion of Nir2 in our HEK293T reporter line (Supp. Fig 1 A) resulted in a decreased rate in the recovery of plasma membrane PIP_2_ levels following stimulation with carbachol (Fig 1D, E).

To measure the effect of Li treatment on PIP_2_ resynthesis, we incubated the reporter line with 1 mM Li; this concentration of LiCl was used in our experiments since the therapeutic range for Li medication in BD patients was 0.75 to 1.5 mEq/L (Francis et al., 2004). Following Li treatment, we stimulated the cells with carbachol and compared the dynamics of plasma membrane PIP_2_ levels with that in non-Li treated cells. We found that in Li treated cells the rate with which PIP_2_ levels at the plasma membrane recovered to pre-stimulation levels was significantly slower than in untreated controls (Fig 1F, G). These findings strongly suggest that treatment with Li results in a reduction in the rate of PIP_2_ resynthesis following PLC stimulation.

In an alternative approach, we quantified total cellular PIP_2_ mass using liquid chromatography coupled with tandem mass spectrometry (LC-MS/MS) at three distinct time points-corresponding to the basal state, one-minute post-stimulation of PLC via carbachol addition (hydrolysis of PIP_2_) state and post-stimulation (recovery of PIP_2_) state. From these experiments, the time for recovery of PIP_2_ (to pre-stimulation levels) was standardized at 15 minutes post stimulation with carbachol (Fig 1H). Using these conditions, we compared the total PIP_2_ mass between untreated cells and those treated with Li. We found that the mass of PIP_2_ during the recovery phase was lower in Li treated cells compared to controls (Fig 1I); likewise, the level of phosphatidylinositol phosphate (PIP) in Li treated cells was also lower during the recovery phase (Fig 1J). Thus, Li treatment lowers the recovery of PIP and PIP_2_ levels following PLC activation.

### Chronic Li treatment reduces PLC dependent intracellular Ca^2+^ mobilization

A key outcome of agonist triggered, PLC mediated PIP_2_ hydrolysis is intracellular Ca^2+^ signalling (Berridge, 2009). Since we noted a reduction in the turnover of PIP_2_ following Li treatment, we tested the impact of Li treatment on intracellular Ca^2+^ levels [Ca^2+^]_i_. Using the control-M1 cell line, we monitored [Ca^2+^]_i_ under both resting conditions as well as stimulation with carbachol, an agonist of m1AchR. Cells were incubated in 1 mM Li prior to the experiment: basal [Ca^2+^]_i_ were no different between treated and untreated cells (Fig 2A). Stimulation of untreated cells with carbachol evoked a rise in [Ca^2+^]_i_ which then decayed back to resting levels (Fig 2B). In cells treated with Li, the carbachol induced rise in [Ca^2+^]_i_ was reduced compared to that seen in untreated cells (Fig 2C). Receptor activated PLC mediated Ca ^2+^ signalling is a biphasic process with two components, an initial release of Ca^2+^ from intracellular stores followed by store-operated influx of Ca^2+^ into the cell from the extracellular medium (Prakriya and Lewis, 2015). We performed experiments to separately monitor each of these components. During the initial phase of the assay, carbachol stimulation was performed in the presence of zero extracellular Ca^2+^, thus selectively monitoring release of Ca^2+^ from intracellular stores (Fig 2D, E). Following this, the extracellular solution was supplemented with Ca^2+^ and the influx of Ca^2+^ into the cytoplasm was monitored (Fig 2D, F). This analysis revealed that in Li treated cells both intracellular Ca^2+^ release (Fig 2E) as well as the subsequent store operated Ca^2+^ influx into cells (Fig 2F) was reduced compared to Li untreated controls. This reduction in receptor activated release of Ca^2+^ from intracellular stores was not due to a reduction in the size of intracellular Ca^2+^ stores as emptying of stores by the application of thapsigargin resulted in equivalent rises in [Ca^2+^]_i_ in Li treated cells and untreated controls (Fig 2G, H).

**Figure 2:**
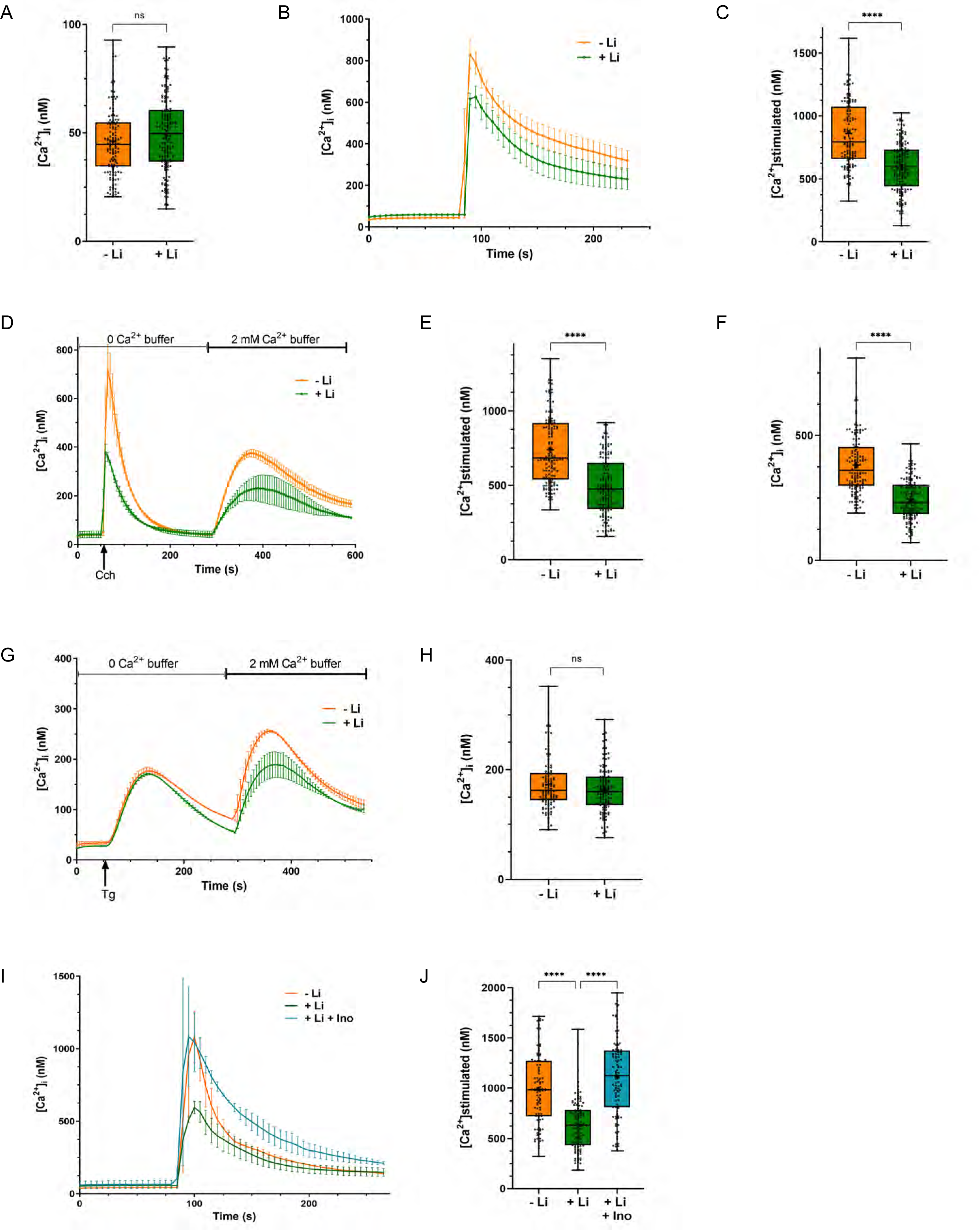
Chronic Lithium treatment reduces PLC dependent intracellular Ca^2+^ mobilization. (A) Basal [Ca^2+^]_i_, calculated in nM using Grynkiewicz equation, is unchanged in control-M1 cells due to lithium treatment (untreated cells-represented by orange, n = 133; cells treated with Li -represented by green line, n =156). (B-C) PLC dependent calcium mobilization is decreased in lithium treated cells. Quantification is done from multiple experiments, each with replicates (untreated cells (orange), n =133; cells treated with Li (green), n =156). (D-F) In zero Ca^2+^ buffer, agonist dependent calcium mobilization is reduced, and subsequent Store Operated Calcium Entry (SOCE) is also decreased in lithium treated cells. Quantification: untreated cells (orange), n =145; cells treated with Li (green), n =146. (G-H) Intracellular store measured by Tg, is unaltered by lithium treatment. Quantification: untreated cells (orange), n =114; cells treated with Li (green), n =118. (I-J) Inositol supplementation reversed the lithium mediated decrease in intracellular Ca^2+^ release. Quantification: untreated cells (orange), n =98; cells treated with Li (green), n =114, cells treated with Li and high inositol (blue), n =104. [Statistical tests: (C, E, F) Student’s unpaired t test. *p value < 0.05; **p value <0.01; ***p value < 0.001; ****p value < 0.0001 and (J) one-way ANOVA with post hoc Tukey’s multiple pairwise comparisons. *p value < 0.05; **p value <0.01; ***p value < 0.001; ****p value < 0.0001].

According to the “inositol depletion hypothesis” (Berridge et al., 1989), Li can restrict the available cytoplasmic inositol pool available for phosphatidylinositol (PI) (and thence PIP_2_) resynthesis by inhibiting IMPase-thereby, lowering down the synthesis of the pool of PIP_2_ that is required for PLC activity. This cytoplasmic inositol pool can also be replenished by transport of inositol from the extracellular medium via plasma membrane inositol transporters [summarized in (Balla, 2013)]. This implies that if a deficit in the available pool of cytosolic inositol is responsible for the attenuated calcium signalling in Li treated cells, this might be rescued by supplementation of the extracellular medium with inositol. We grew HEK293T cells in an inositol rich DMEM media (inositol concentration is ca. 28 mg/litre; which is similar to the inositol concentration in the cerebrospinal fluid (Nixon, 1955; Shetty et al., 1996; Swahn, 1985)). These cells were subjected to Li treatment and agonist induced Ca^2+^ influx was quantified. As controls in this experiment, we used Li treated control-M1 cells grown in DMEM not supplemented with inositol. As in previous experiments, we found that when cells were grown in DMEM, Li treatment resulted in a reduction of agonist induced Ca^2+^ influx (Fig 2I, J). However, when cells grown in inositol supplemented DMEM were treated with Li, the Li induced reduction in Ca^2+^ influx was rescued compared to that seen in cells grown under equivalent Li treatment but in DMEM not supplemented with inositol (Fig 2I, J), compared to Li non-treated cells.

### IMPA1 is required for agonist mediated calcium signalling and PIP_2_ resynthesis

IMPase, the enzyme that dephosphorylates inositol monophosphate to generate myo-inositol has been proposed as a direct target of the action of Li, inhibiting its activity (Hallcher and Sherman, 1980; Saudek, 1996). There are two genes encoding IMPase in the human genome *IMPA1* and *IMPA2* of which *IMPA1* is the predominant isoform expressed in HEK293T cells. To test the requirement of *IMPA1*, we generated a HEK293T cell line in which *IMPA1* was deleted by CRISPR/Cas9 genome editing (Supp. Fig 3A); m1AchR was expressed in these cells by lentiviral transduction; these cells will be denoted as IMPA1^-/-^-M1. Western blot analysis of IMPA1^-/-^-M1 showed the complete absence of the IMPA1 protein (Fig 3A), IMPA2 transcript levels were not altered (Supp. Fig 3B) and the levels of m1AchR protein was comparable between control-M1 cells and IMPA1^-/-^-M1 cells (Supp. Fig 3C, D). We compared the response of IMPA1^-/-^-M1 cells to control-M1 cells. When stimulated with carbachol, IMPA1^-/-^-M1 cells showed a reduced Ca^2+^ influx response that could be rescued by reconstitution of the cells with the IMPA1 cDNA (Fig 3 A-C). Further analysis under conditions of zero extracellular Ca^2+^ revealed that the release of Ca^2+^ from intracellular stores was reduced in in IMPA1^-/-^-M1 cells compared to control cells (Fig 3D, E) with an intact *IMPA1* gene. Lastly, we found that following carbachol stimulation, the rate at which PIP_2_ levels were restored at the plasma membrane was slower in IMPA1^-/-^-M1 and this could be rescued by reconstitution with a wild type IMPA1 transgene (Fig 3F).

**Figure 3:**
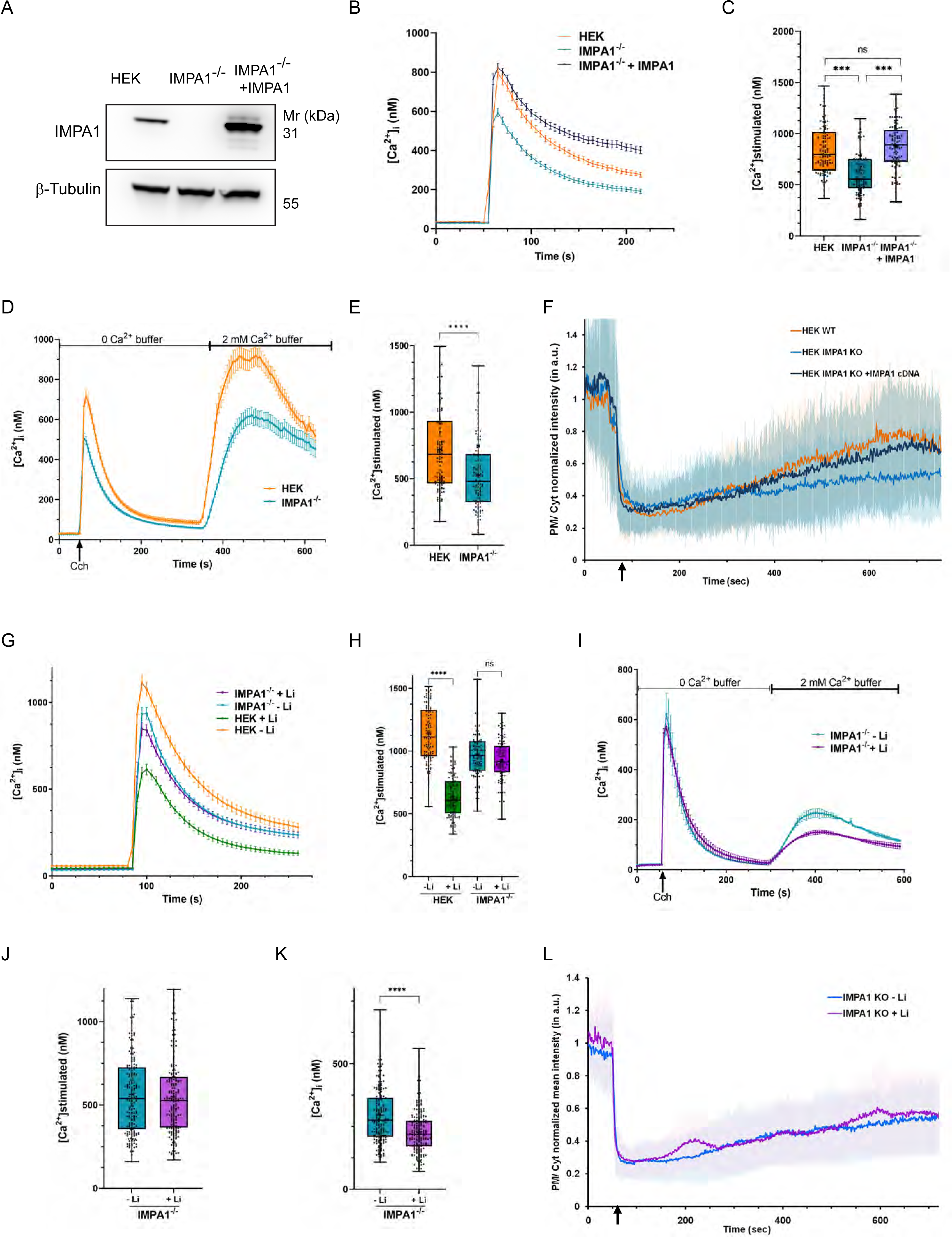
IMPA1 is required for the effect of Li on PIP_2_ resynthesis and agonist mediated Ca^2+^ signalling. (A) Western blot showing IMPA1 expression in IMPA1^-/-^ cells and IMPA1^-/-^ cells transduced by *IMPA1* cDNA. (B, C) PLC dependent calcium mobilization is decreased in IMPA1^-/-^ cells; *IMPA1* expression via lentiviral transduction of cDNA in HEK IMPA1^-/-^ cells rescued the decreased intracellular Ca^2+^ release. Quantification: untreated HEK cells (orange), n =155; HEK IMPA1^-/-^ cells (light blue), n =143; HEK IMPA1^-/-^ cells + IMPA1 cDNA (dark blue), n =162. (D, E) In zero Ca^2+^ buffer, agonist dependent calcium mobilization is decreased in HEK IMPA1^-/-^ cells, compared to HEK cells. Quantification: HEK cells (orange), n =121; HEK IMPA1^-/-^ (blue), n =120. (F) PIP_2_ turnover post PLC stimulation in HEK cells compared to HEK IMPA1^-/-^ cells; *IMPA1* expression via lentiviral transduction of cDNA in HEK IMPA1^-/-^ cells rescued the delay in PIP_2_ turnover. Quantification: untreated HEK cells (orange), n =51; HEK IMPA1^-/-^ cells (light blue), n =50; HEK IMPA1^-/-^ cells + IMPA1 cDNA (dark blue), n =44. (G, H) Cch mediated calcium release in these IMPA1^-/-^ cells is not altered by lithium. Quantification: untreated HEK cells (orange), n =126; HEK cells treated with Li (green), n =127, HEK IMPA1^-/-^ cells without Li treatment (blue), n =132; HEK IMPA1^-/-^ cells treated with Li (purple), n =112. (I-K) In zero Ca^2+^ buffer, agonist dependent calcium mobilization is unaltered, but subsequent Store Operated Calcium Entry (SOCE) is decreased in lithium treated IMPA1^-/-^ cells. Quantification: HEK IMPA1^-/-^ cells without Li treatment (blue), n =179; HEK IMPA1^-/-^ cells treated with Li (purple), n =193. (L) Lithium does not alter the rate of regeneration of PIP_2_ at the plasma membrane in the IMPA1^-/-^ cells, as monitored by the changes in the translocation of the PH-PLCδ::GFP probe. Mean ± SD is plotted from four experiments, each performed in replicates (HEK IMPA1^-/-^ cells without Li treatment-blue line, n =50; HEK IMPA1^-/-^ cells treated with Li-purple line, n =47). [Statistical tests: (C, H) one-way ANOVA with post hoc Tukey’s multiple pairwise comparison. *p value < 0.05; **p value <0.01; ***p value < 0.001; ****p value < 0.0001 and (E, J, K) Student’s unpaired t test. *p value < 0.05; **p value <0.01; ***p value < 0.001; ****p value < 0.0001].

### IMPA1 is required for the effect of Li on receptor activated Ca^2+^ influx and PIP_2_ resynthesis

We tested the effect of Li treatment on IMPA1^-/-^-M1 compared to untreated IMPA1^-/-^-M1 cells. We analysed receptor activated Ca^2+^ influx, following Li treatment; this was not altered compared to untreated IMPA1^-/-^-M1 cells (Fig 3G, H); in the same experiment, treatment of control-M1 cells with Li resulted in a big reduction in agonist activated Ca^2+^ influx. We also analyzed the two phases of receptor activated Ca^2+^ influx and found that while release of [Ca^2+^]_i_ from stores was not affected by Li treatment in IMPA^-/-^-M1 cells, the reduction of SOCE previously noted on Li treatment in control-M1 cells was still present (Fig 3I-K). We also found that the rate of regeneration of PIP_2_ at the plasma membrane was unchanged in IMPA1^-/-^-M1 cells subjected to Li treatment, compared to untreated IMPA1^-/-^-M1 cells (Fig 3L, Supp. Fig 3E). These observations suggest that the ability of Li to inhibit Ca^2+^ influx and slow PIP_2_ resynthesis following PLC activation requires an intact IMPA1.

### Li reduces excitability in human cortical neurons

To test the relevance of our findings to human cortical neurons, changes in whose activity presumably underlies the therapeutic effect of Li in bipolar disorder patients, we generated human forebrain cortical neurons from induced pluripotent stem cells (iPSC) *in vitro* (Sharma et al., 2020). We differentiated forebrain cortical neurons from neural stem cells (NSC) derived from an Indian control iPSC line, D149 (Fig 4A) (Iyer et al., 2018). The generated NSCs expressed previously described markers characteristic of NSC such as Nestin, SOX1, SOX2 and PAX6 (Fig 4B) and when differentiated, expressed the neuronal marker MAP2; a proportion of cells in the culture expressed the glial marker GFAP (Fig 4C). When these NSCs were differentiated into cortical neurons, they exhibited spontaneous [Ca^2+^]_i_ elevations, referred to as transients, that increase as a function of age *in vitro* (Akhtar et al., 2022; Sharma et al., 2020). We measured somatic [Ca^2+^]_i_ transients from such neuronal cultures at 45 *days in vitro* (DIV 45) and quantified their frequency. We observed that the frequency of [Ca^2+^]_i_ transients was decreased following Li treatment (1 mM) for 10 days (Fig 4D, E). We also measured changes in [Ca^2+^]_i_ following stimulation with carbachol in DIV45 cortical neurons treated with Li. In our cultures carbachol stimulation results in a rise in [Ca^2+^]_i_ that peaks before gradually decaying towards the baseline; in Li treated cultures the peak of this response was reduced (Fig 4F, G). We also performed experiments to measure Ca^2+^ mobilization from internal stores; these revealed that carbachol activated release of Ca^2+^ in Li treated neurons was reduced compared to those that were untreated (Fig 4H-J). By contrast there was only a modest difference in thapsigargin mediated rise in [Ca^2+^]_i_ between Li treated and untreated cortical neurons (Supp. Fig 4A-C). These observations suggest that both neuronal excitability as well as PLC mediated Ca^2+^ signalling is attenuated in human cortical neurons following Li treatment.

**Figure 4:**
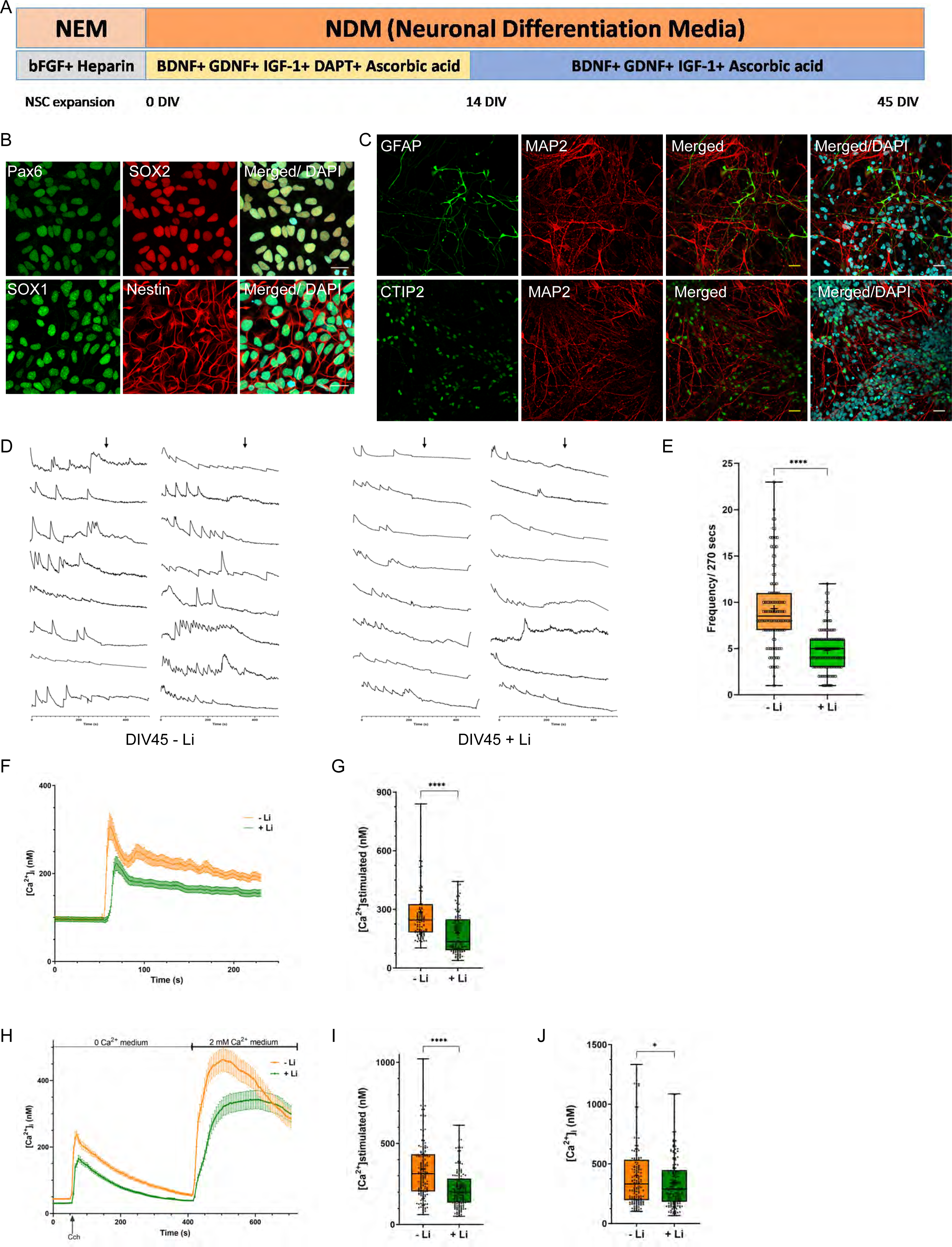
Lithium reduces excitability in human cortical neurons. (A) Overview of expansion and differentiation process of the D149 lineage Neural Stem Cells (NSCs) and the cortical neurons. (B) Characterization of NSCs via immunocytochemistry of known markers like Nestin, Sox2, Sox1 and Pax6. (C) The mature cortical neurons are characterized via immunocytochemistry of markers like MAP2, CTIP2 and GFAP (Astrocytes marker) [Scale Bar – 25 μM]. (D) [Ca^2+^]_i_ traces from individual cells (soma; Y-axis shows ΔF and X-axis is time in seconds. The baseline was recorded for 270 secs followed by addition of 10 μM Tetrodotoxin (TTX) (as indicated by the arrows) to block Ca^2+^ transients. Transients are higher in frequency in the neuronal cultures without lithium treatment, compared to the treated cultures. The number of spikes/ 270 secs are counted from individual soma and plotted, each dot representing events from a single soma. (E) Data is plotted from multiple experiments, each done with biological replicates (untreated neurons (orange), n =114; neurons treated with Li (green), n =113). (F-G) Cch dependent calcium mobilization was decreased in lithium treated neuronal cultures (DIV 45). Quantification: untreated neurons (orange), n =97; neurons treated with Li (green), n =106. (H-J) Decrease in the Cch dependent calcium mobilization as well as in Store Operated Calcium Entry (SOCE) cells in lithium treated neuronal cultures (DIV 45). Quantification: untreated neurons (orange), n =154; neurons treated with Li (green), n =156.[Statistical tests: (E,G,I,J) Student’s unpaired t test. *p value < 0.05; **p value <0.01; ***p value < 0.001; ****p value < 0.0001].

### Inhibition of GSK-3β does not phenocopy the effect of Li on human cortical neurons

It has been proposed that the inhibition of GSK-3β by Li underlies its therapeutic effects in BPAD. To test this hypothesis, we treated HEK293T cells with the well-established pharmacological inhibitor of this enzyme-CHIR99021(An et al., 2014). Treatment of control-M1 cells with 10μM CHIR99021 for 24 hrs resulted in a robust elevation of β-catenin levels indicating potent inhibition of GSK-3β under these conditions (Fig 5A). Under these conditions, we found that GSK-3β inhibition did not result in reduced agonist induced Ca^2+^ influx (Fig 5B, C). Likewise, in human iPSC derived forebrain cortical neurons, the frequency of spontaneous Ca^2+^ transients (Fig 5D, E) and carbachol activated Ca^2+^ influx was not altered by GSK-3β inhibition (Fig 5F, G). Lastly, following PLC activation and PIP_2_ depletion, the rate at which PIP_2_ levels at the plasma membrane were restored was not impacted by treatment of cells with the GSK-3β inhibitor (Fig 5H).

**Figure 5:**
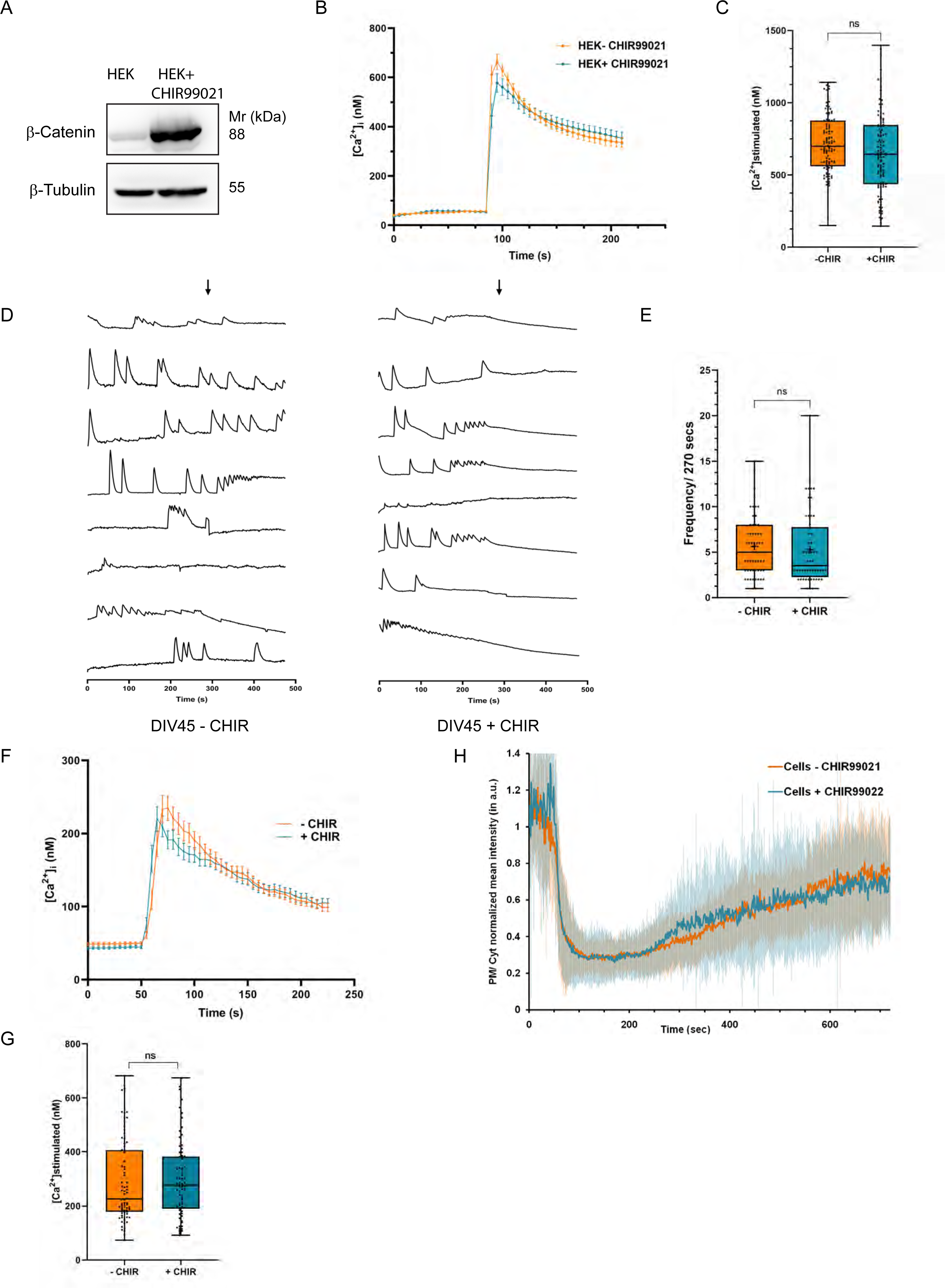
GSK-3β inhibition by CHIR99021 does not affect PLC mediated Ca^2+^ mobilization or PIP_2_ turnover. (A) Western blot showing CHIR99021 (an inhibitor of GSK-3β) treatment led to the high expression of β-Catenin, one of the targets of GSK-3β destruction complex. (B, C) Inhibition of GSK-3β by CHIR99021 did not alter agonist mediated Ca^2+^ release. Quantification: untreated cells (orange), n =116; cells treated with CHIR99021 (blue), n =118. (D) [Ca^2+^]_i_ traces from individual cells (DIV45 neurons) untreated or treated with CHIR99021 (soma; Y-axis shows ΔF and X-axis is time in seconds). The number of spikes/ 270 secs are counted from individual soma and plotted, each dot representing events from a single soma. (E) Quantification-untreated neurons (orange), n =55; neurons treated with CHIR99021 (blue), n =52. (F, G) Cch dependent calcium mobilization was unchanged in CHIR99021 treated neuronal cultures (DIV45). Quantification: untreated neurons (orange), n =76; neurons treated with CHIR99021 (blue), n =74. (H) PIP_2_ turnover post PLC stimulation in HEK cells treated with CHIR99021, compared to untreated cells (untreated cells-orange line, n =31, cells treated with CHIR99021-blue line, n =34). [Statistical tests: (B, C) Student’s unpaired t test. *p value < 0.05; **p value <0.01 ***p value < 0.001; ****p value < 0.0001].

### Li treatment induces transcriptional changes in pathways involved in neuronal excitability

To understand the molecular mechanisms underlying the reduced excitability following Li treatment in human iPSC derived cortical neurons, we performed a transcriptomic analysis comparing untreated DIV45 neurons with Li treated DIV45 neurons (Fig 6A). Differentially expressed genes were obtained using DeSeq2; a cut-off of log_2_FC change greater than 0.2 was used along with a p-value and FDR significance at less than 0.05 were considered. Using these criteria, 335 upregulated and 361 downregulated genes were obtained (Supplementary Table 1 S1, S2). Gene Ontology (GO) analysis of the KEGG pathway showed substantial enrichment for genes annotated as being involved in calcium signalling or in glutamatergic synapse (Supplementary Table 1 S3). In addition, GO KEGG analysis of the downregulated gene set also showed enrichment for several other categories of neurotransmission and neuromodulation related pathways (cholinergic, GABAergic, dopaminergic, glutamatergic and serotonergic) (Supplementary Table 1 S2) (Fig 6B).

**Figure 6:**
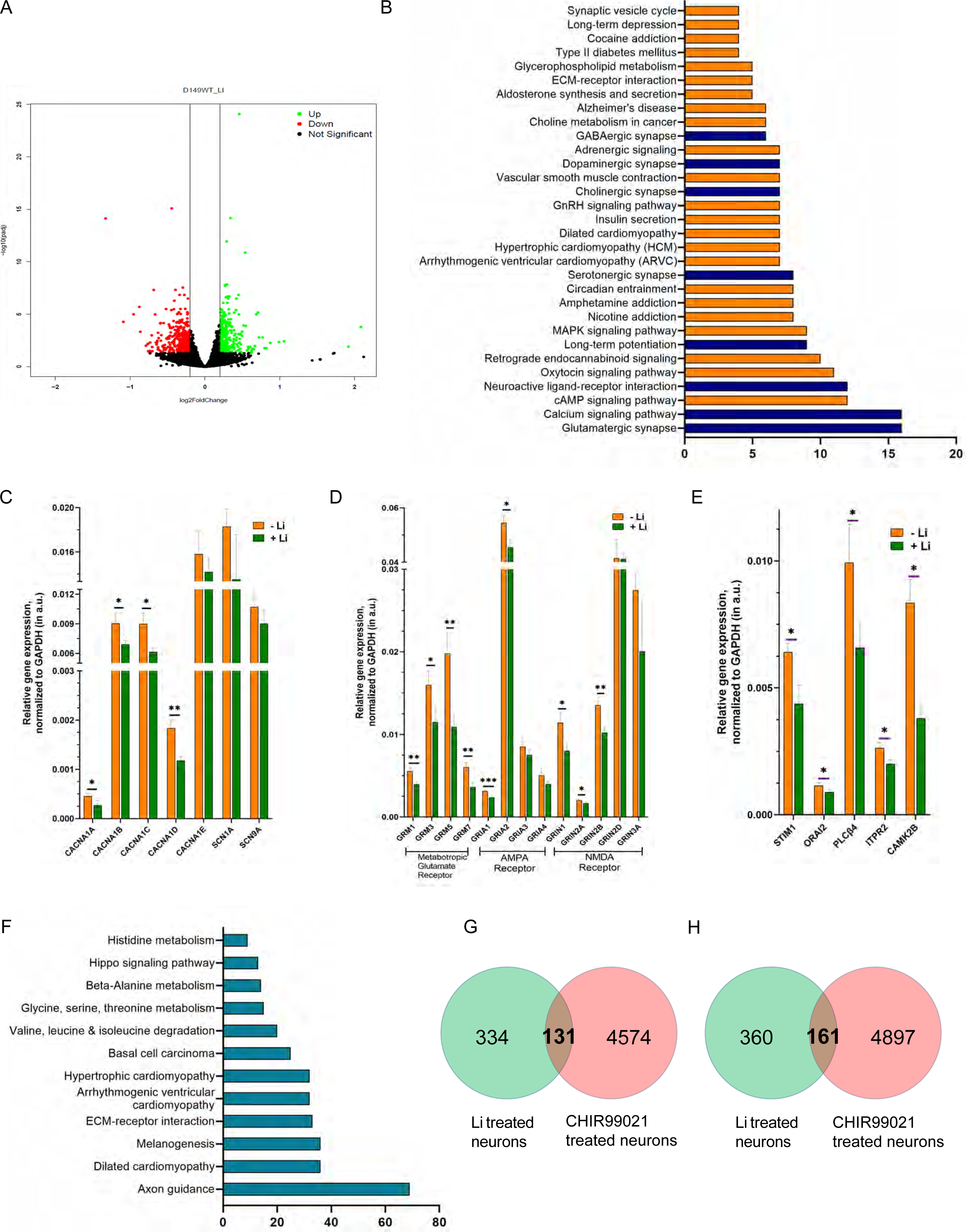
Lithium treatment induces transcriptional changes in pathways involved in neuronal transmission. (A) Volcano plot showing expression of genes altered due to lithium treatment in human cortical neuronal cultures DIV45 (p value <0.05, p adj value <0.01). (B) The GO KEGG pathway enrichment analysis for downregulated genes. (C-E) Quantitative analysis showing several genes related to voltage calcium channels (C), Glutamatergic pathway (D) and calcium homeostasis (E), are downregulated in cortical neurons (DIV45) due to lithium treatment. (F) The GO KEGG pathway enrichment analysis for downregulated genes due to CHIR99021 treatment. (G,H) Distribution of the number of upregulated and downregulated genes in neuronal cultures (DIV45) due to Li treatment and CHIR99021 treatment. [Statistical tests: (C, D, E) Multiple unpaired t test. *p value < 0.05; **p value <0.01; ***p value < 0.001].

Since GO analysis revealed an enrichment in Ca^2+^ signalling genes, we analysed molecular mechanisms through which Li might downregulate neuronal Ca signalling and hence excitability. There are multiple molecular mechanisms by which neuronal calcium signalling is modulated (Berridge, 1998) including voltage gated Ca channels and receptor activated Ca signalling, a mechanism used by several neurotransmitters and neuromodulators. Using the transcriptomic data, we tested the likely contribution of each of these mechanisms to the effects of Li in our model system. Physiological and pharmacological evidence suggests that the [Ca^2+^]_i_ transients seen during neuronal development arise from the activity of voltage gated calcium channels (VGCC) (Rosenberg and Spitzer, 2011) and this also seems to underlie the [Ca^2+^]_i_ transients seen in our experiments when human iPSC are differentiated into cortical neurons (Sharma et al., 2020)[this study]. To understand the mechanism by which Li could reduce the frequency of [Ca^2+^]_i_ transients we studied the expression of genes encoding VGCC. Our transcriptomic analysis suggested that expression of genes encoding several subunits of VGCC might be downregulated (Supplementary Table 1 S3). We experimentally tested the expression of genes for *CACNA1A, CACNA1B, CACNA1C, CAC1A1D* and *CACN1A1E* by qRT-PCR analysis; this showed that the expression of CACNA1A,B,C,D were all downregulated in Li treated DIV45 neurons compared to controls (Fig 6C) whereas CACNA1E was unchanged. In the same analysis, there was no downregulation in transcripts for *SCN1A (Nav1.1)* or *SCN9A (Nav7.1)* that encode the brain enriched, α-subunit of voltage gated sodium channels, involved in plasma membrane depolarization that trigger VGCC dependent [Ca^2+^]_i_ transients (Fig 6C).

Glutamate is a key excitatory neurotransmitter in the human central nervous system, exerting its effects through both metabotropic and ionotropic mechanisms (Reiner and Levitz, 2018) and changes in [Ca^2+^]_i_ is a key mechanism in glutamate signalling in the human brain. Our qRT-PCR analysis of genes encoding key components in the glutamate signalling system in Li treated neurons revealed that transcript levels for metabotropic glutamate receptors (*GRM1, GRM3, GRM5, GRM7*) and the ionotropic glutamate receptors (*GRIA1, GRIA2, GRIN1, GRIN2A, GRIN2B*) were downregulated (Fig 6D).

Many neurotransmitters and modulators transduce signal via G-protein coupled, PLC mediated store-operated Ca^2+^ influx (Moccia et al., 2015). Key molecular components of this pathway include the IP_3_ receptor, ER Ca^2+^ sensor STIM and the Ca influx channel ORAI; we found that transcript levels of *PLCb4, ITPR2, STIM1* and *ORAI2* were all downregulated in Li treated neurons compared to controls (Fig 6E).

### Li and GSK-3β inhibition induce distinct transcriptional responses

We also examined the transcriptional response of human iPSC derived forebrain cortical neurons to treatment with 10μM CHIR99021 for 24 hrs. RNA-seq analysis revealed large changes in gene expression with ca. 5000 genes each being up or downregulated (Supp. Fig 6C). GO analysis revealed a broad range of pathways including cardiomyopathy, amino acid metabolism, and extracellular matrix interactions (Fig 6F). Strikingly, there was no enrichment of pathways related to synaptic transmission or calcium signalling; the only brain related pathway that was enriched was axon guidance and the identity of the genes picked up suggested ECM function. We overlapped the set of genes up or downregulated in Li and GSK-3β inhibition transcriptomes to assess the extent of overlap between the two gene sets and found that both in the domain of up and downregulated genes (Fig 6G, H), there was at best a 3% overlap in terms of altered genes. These findings suggest that at the level of transcriptional changes, the mechanism of action of Li and GSK-3β is distinctive.

### Mechanisms of Li induced transcriptional changes

To understand the mechanisms underlying Li induced transcriptome changes we analysed the likely transcriptional control mechanisms. For this we used the htFtarget database that has information on all human transcription factors (TFs) and the genes they control, tissue specific TF-target information, data on TFs for non-coding RNA and information on co-regulation for a target gene and its TFs (Zhang et al., 2020). TFs controlling genes differentially expressed in Li treated neurons (DIV 45) were identified from the htFtarget database. Out of 650 up regulated genes in Li treated neurons, 587 were identified as associated with one or more TFs in this database. Likewise, out of 525 downregulated genes, 333 genes were linked to a TF. 485 and 323 unique TFs were identified that control the upregulated and downregulated genes respectively. We then further looked for transcription factors that control differentially expressed genes that are involved in calcium signalling (Supp. Table1 S3) (Supp. Fig 6A). Of the upregulated genes, only RYR receptor was associated with three different transcription factors (CTCF, SPI1 and USF1) that could control its expression. On the other hand, several genes involved in calcium signalling that were downregulated were controlled by transcription factors. Expression of *CACNA1C, CACNA1D, CAMK2B, CAMK4, CHRM3, GRIN1, ITPR2, RYR3, SCN9A* and *SLC2A13* genes is regulated by transcription factors. Of these *ITPR2* and *CAMK2B* are regulated by 10 different TFs, while *CACNA1D, CHRM3, RYR3, SCN9A and SLC2A13* are regulated by single transcription factor (Supp. Fig 6B). This suggests that some of the genes that are differentially expressed in Li treated neurons are under transcriptional control which can be a direct or indirect effect of Li treatment.

## Discussion

Although Li is a monovalent ion with a remarkable therapeutic effect in the management of BPAD, there has been a lack of clear understanding on the mechanisms of action of Li in brain cells. Although many molecular targets of Li have been described in the literature, their role in mediating the action of Li in human brain cells has not been directly tested.

One of the earliest cellular effects described for Li was its ability to slow phosphoinositide turnover in neural cells (Berridge et al., 1982) and it has been proposed that the consequent slowing of neurotransmitter activated neuronal excitability may underlie the effectiveness of the compound in managing BPAD. Since Li inhibits the enzyme IMPA1, one long-standing proposal is that the application of Li to cells disrupts the receptor activated PIP_2_ cycle and its downstream effect on [Ca^2+^]_i_ signalling leading to reduced neuronal excitability. However, there has not been a direct test of the model that the ability of Li to modulate inositol turnover and [Ca^2+^]_i_ signalling *in vivo* requires the *IMPA1* gene product. In this study, we found that in the application of Li to human cells reduced [Ca^2+^]_i_ signalling following PLC activation. We find that this is underpinned by a reduction in the release of Ca^2+^ from intracellular stores. During PLC activation, the principal mechanism by which Ca^2+^ is released is via the IP_3_ receptor; however, we found that neither the transcript levels for this gene (Supp. Fig 2C) nor the size of stores itself was altered by Li treatment (Fig 2G, H). Thus, it seems most likely that Li treatment affects IP_3_ receptor mediated Ca^2+^ release by altering IP_3_ production during receptor mediated PLC stimulation, a process that requires adequate levels of the substrate for PLC, PIP_2_. Our finding that Li treatment slows synthesis of PIP_2_, the substrate from which PLC produces IP_3_ likely provides an explanation for the reduced intracellular Ca^2+^ release phenotype seen in Li treated cells following PLC stimulation.

If Li exerts its effects in neurons via inhibition of IMPA1, a prediction is that cells lacking IMPA1 might be unresponsive to Li. We generated a cell line in which IMPA1 was deleted and found that in the absence of IMPA1, Li was unable to exert its effects on both [Ca^2+^]_i_ signalling and PIP_2_ resynthesis during receptor activated PLC signalling. These findings imply that an intact IMPA1 is required for the action of Li on these processes. Consistent with this idea, we noted that receptor activated [Ca^2+^]_i_ signalling (Fig 3D, E) and the rate of PIP_2_ resynthesis (Fig 3F) following PLC activation was reduced in IMPA1^-/-^ cells without Li treatment. Therefore, this study provides compelling evidence for the link between the inhibition of IMPA1 by Li leading to reduced PIP_2_ synthesis and thence to reduced neurotransmitter activated, PLC mediated Ca^2+^ signalling. Although we generated IMPA1^-/-^ NSC, we were unable to differentiate these into cortical neurons as NSC of this genotype undergo cell death soon after the initiation of differentiation. This finding is consistent with a previous report that stem cell lines from a human patient with an IMPA1 mutation could not be differentiated into cortical neurons (Figueiredo et al., 2021).

How does IMPA1 regulate PIP_2_ synthesis and [Ca^2+^]_i_ signalling? IMPA1 dephosphorylates inositol 1-phosphate to generate inositol. Hence biological effects resulting from Li inhibition of IMPA1 could arise either from an accumulation of the substrate, inositol 1-phosphate or a deficiency of the product, inositol. Indeed, Li treatment in rodent models has shown both an elevation of inositol 1-phosphate levels as well as a modest reduction in inositol levels (Sade et al., 2016; Sherman et al., 1981). In our analysis, we found that the lowered [Ca^2+^]_i_ influx following receptor activation in human cells could be rescued by supplementation of the extracellular medium with inositol. This observation implies that the [Ca^2+^]_i_ signalling defect arising from Li inhibition of IMPA1 is likely to be a consequence of inositol depletion rather than an accumulation of the substrate inositol 1-phosphate. During the PLC activated PIP_2_ cycle, inositol generated by the action of IMPA1 is condensed with cytidine diphosphate diacylglycerol to form phosphatidylinositol (Lykidis et al., 1997) which is then sequentially phosphorylated to generate PIP_2_. In this study, we found that following PLC activation, the resynthesis of PIP_2_ was also slowed by treatment of wild type cells with Li. This observation is consistent with our finding that PIP_2_ resynthesis and [Ca^2+^]_i_ signalling were both reduced in IMPA1^-/-^ cells. Together, our data provide compelling evidence that Li treatment results in inhibition of IMPA1, a depletion of the inositol pool required for PIP_2_ resynthesis during PLC signalling and hence reduced activity of neurons during neurotransmitter activated synaptic transmission.

An alternative and widely discussed molecular target of Li is GSK-3β. However, in this study, we found that in the hiPSC derived cortical neuron cultures, inhibition of GSK-3β did not phenocopy the physiological effects of Li treatment on neuronal excitability and PLC signalling. Further, RNA seq analysis revealed minimal overlap in the transcriptome changes induced by Li treatment and GSK-3β inhibition. Thus, it seems, in the hiPSC model system, Li is unlikely to influence neuronal excitability via GSK-3β inhibition.

When Li is used to treat BPAD in human patients, presumably it acts by reducing the excitability of neurons in the cerebral cortex. In this study, we tested the effect of Li treatment on the physiology of human forebrain cortical neuronal cultures differentiated from iPSC. Consistent with previous reports (Mertens et al., 2015), we found that treatment of human forebrain cortical neurons with therapeutically relevant concentrations of Li reduced the frequency of [Ca^2+^]_i_ transients that are underpinned by voltage gated Ca^2+^ channel (VGCC) activity (Sharma et al., 2020). Since acute application of Li does not affect [Ca^2+^]_i_ transients (Supp. Fig 4D,E), it seems unlikely that Li affects neuronal excitability by directly inhibiting the VGCC. However, VGCC mediated [Ca^2+^]_i_ transients are activated by neuronal action potentials which themselves are triggered following activation of post-synaptic G-protein coupled receptors that bind neurotransmitters. Several of these receptors (mGluR1, mGluR5, mAchR and 5HT-2A) when bound to their respective neurotransmitter ligand, use PLC activation as part of their signalling mechanism. Thus, Li could influence neuronal excitability by modulating signalling through such PLC linked receptors. In support of this hypothesis, in this study we found that in human forebrain cortical neurons, in addition to reducing [Ca^2+^]_i_ transients, Li application diminished the elevations of [Ca^2+^]_i_ triggered by application of the mAchR ligand carbachol (Fig 4F-I). Thus, a key mechanism by which Li modulates neuronal excitability may be through downregulation of neurotransmitter signalling of PLC linked GPCRs.

Given the extended timeframe over which Li exerts its therapeutic effect it has been proposed that in addition to directly impacting cellular biochemistry, transcriptional mechanisms may play a role in its biological effects. SOCE that occurs downstream of receptor activated PLC signalling has been shown to regulate transcription in neurons (reviewed in (Mitra and Hasan, 2022)). Our transcriptomic analysis of human forebrain cortical neurons revealed that Li treatment on a clinically relevant time and concentration range results in a large transcriptional response. Likewise, a comparison of Li induced differential gene expression (this study) to that elicited by genetic inhibition of SOCE (Dhanya and Hasan, 2021; Gopurappilly et al., 2018) also revealed only a modest overlap of genes that were differentially regulated (Supp Fig 4F, G). Therefore, it is likely that Li exerts transcriptional changes in neurons independent of its effects on SOCE.

Our analysis of the Li induced transcriptome changes in human forebrain cortical neurons however provide an insight into the mechanisms by which Li might mediate its effects both in cultured forebrain cortical neurons and its clinical effects. We found that the levels of three of the four subunits of VGCC were downregulated in Li treated cells and this could explain in part the ability of Li to reduce [Ca^2+^]_i_ transients in forebrain cortical neurons in culture. A key feature of mania in BPAD patients is heightened neural activity manifested behaviorally, EEG and fMRI studies and presumably Li exerts its therapeutic effect through downregulating this neural activity. Glutamate is the principle excitatory neurotransmitter in the brain and consistent with this we found that transcripts for multiple glutamate receptor subtypes were downregulated in forebrain cortical neurons treated with Li. These included multiple subunits of the metabotropic, AMPA and NMDA subtypes of the glutamate receptor family. Downregulation of these will likely contribute to reducing neural activity following Li treatment manifest clinically as behavioral improvement. Overall, this study demonstrates that the effects of Li on neuronal Ca^2+^ signalling are mediated through an IMPA dependent step and involves a large transcriptional response that downregulates molecular processes that are required for neuronal excitability. The link between Li and transcriptional responses are unclear although studies in yeast have shown that treatment with Li is associated with changes in the transcriptome that also includes changes in inositol biosynthesis.

## Acknowledgements

This work was supported by the Department of Atomic Energy, Government of India, under Project Identification No. RTI 4006, the Department of Biotechnology, Government of India through the Accelerator Program for Discovery in Brain Disorders (BT/PR17316/MED/31/326/2015), the Pratiksha Trust and a Wellcome-DBT India Alliance Senior Fellowship to PR (IA/S/14/2/501540). We thank the NCBS Imaging, Mass spectrometry, Genomics, High performance computing, Stem cell and Biosafety facilities for support.

## Supplementary Table legends

### Supplementary Table 1

**Sheet S1: List of upregulated genes in Li treated neurons DIV45.**

The table shows Gene ID (ENSEMBL ID), Gene names, base Mean, log2 fold change, p value, p adjusted value and gene description (as derived from Deseq2 analysis).

**Sheet S2: List of downregulated genes in Li treated neurons DIV45.**

The table shows Gene ID (ENSEMBL ID), Gene names, base Mean, log2 fold change, p value, p adjusted value and gene description (as derived from Deseq2 analysis).

**Sheet S3: List of genes involved in calcium signalling.**

The table shows Gene ID (ENSEMBL ID), Gene names, base Mean, log2 fold change, p value, p adjusted value and gene description (as derived from Deseq2 analysis).

**Sheet S4: GO pathway analysis for the upregulated genes in Li treated neurons DIV45.**

The table shows GO-Category, GO-Term, Count, Percentage of genes, p value, Gene Ids (ENSEMBL IDs), Fold Enrichment and its significance (as derived from DAVID GO analysis).

**Sheet S5: GO pathway analysis for the downregulated genes in Li treated neurons DIV45.**

The table shows GO-Category, GO-Term, Count, Percentage of genes, p value, Gene Ids (ENSEMBL IDs), Fold Enrichment and its significance (as derived from DAVID GO analysis).

### Supplementary Table 2

List of primers used for qRT-PCR.

The table shows the gene names and the primer sequence in 5’ to 3’ orientation.

