## supplementary data for "IMPA1 dependent regulation of plasma membrane phosphatidylinositol 4,5-bisphosphate turnover and calcium signalling by lithium"

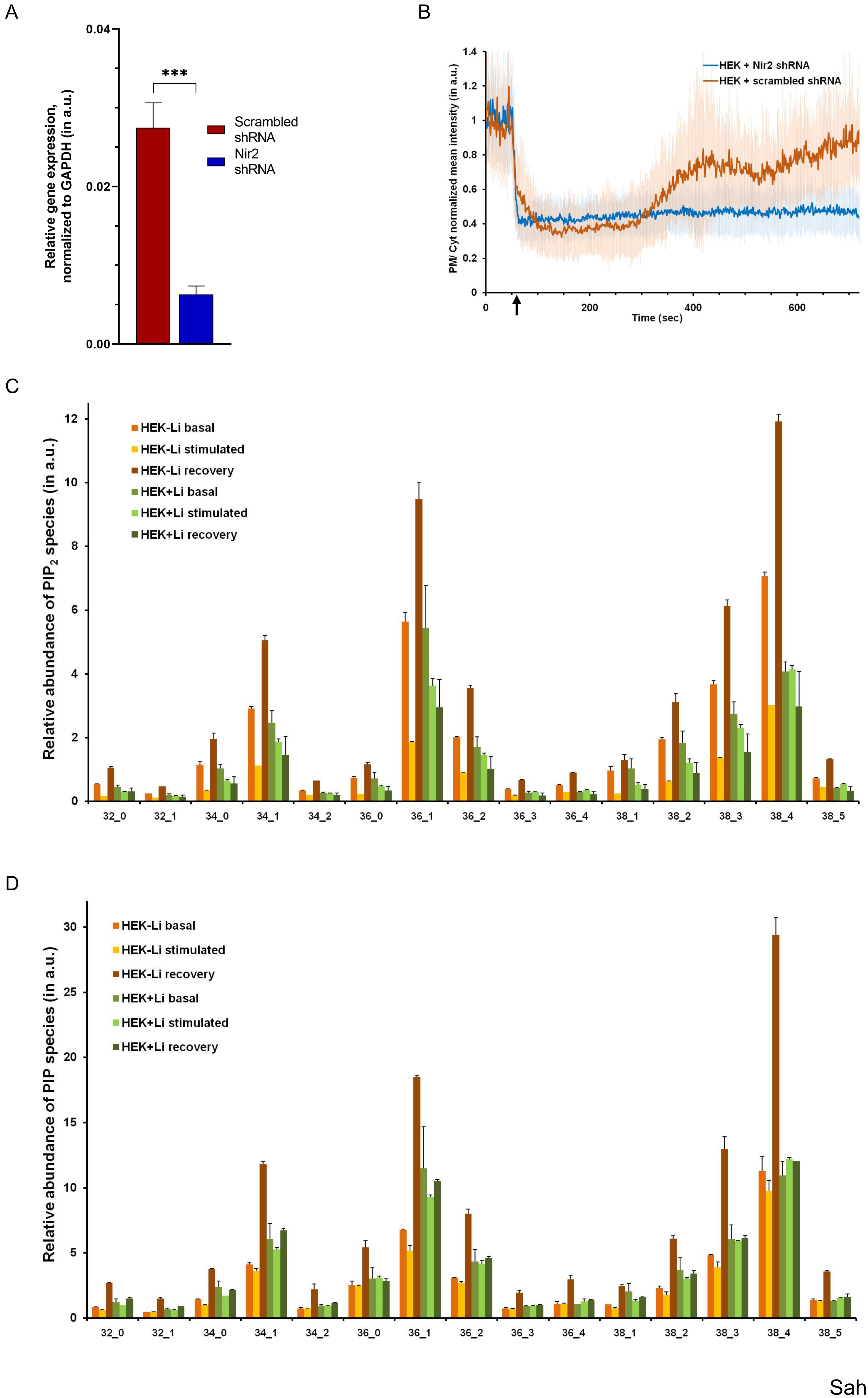


Saha Supp Figure 1

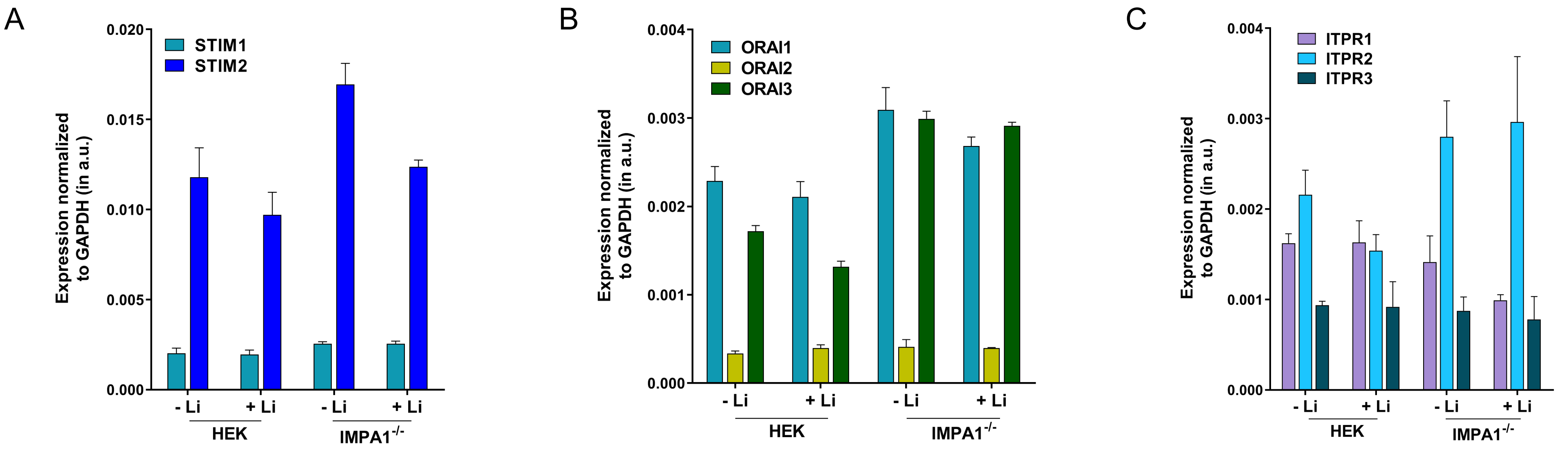


Saha Supp Figure 2

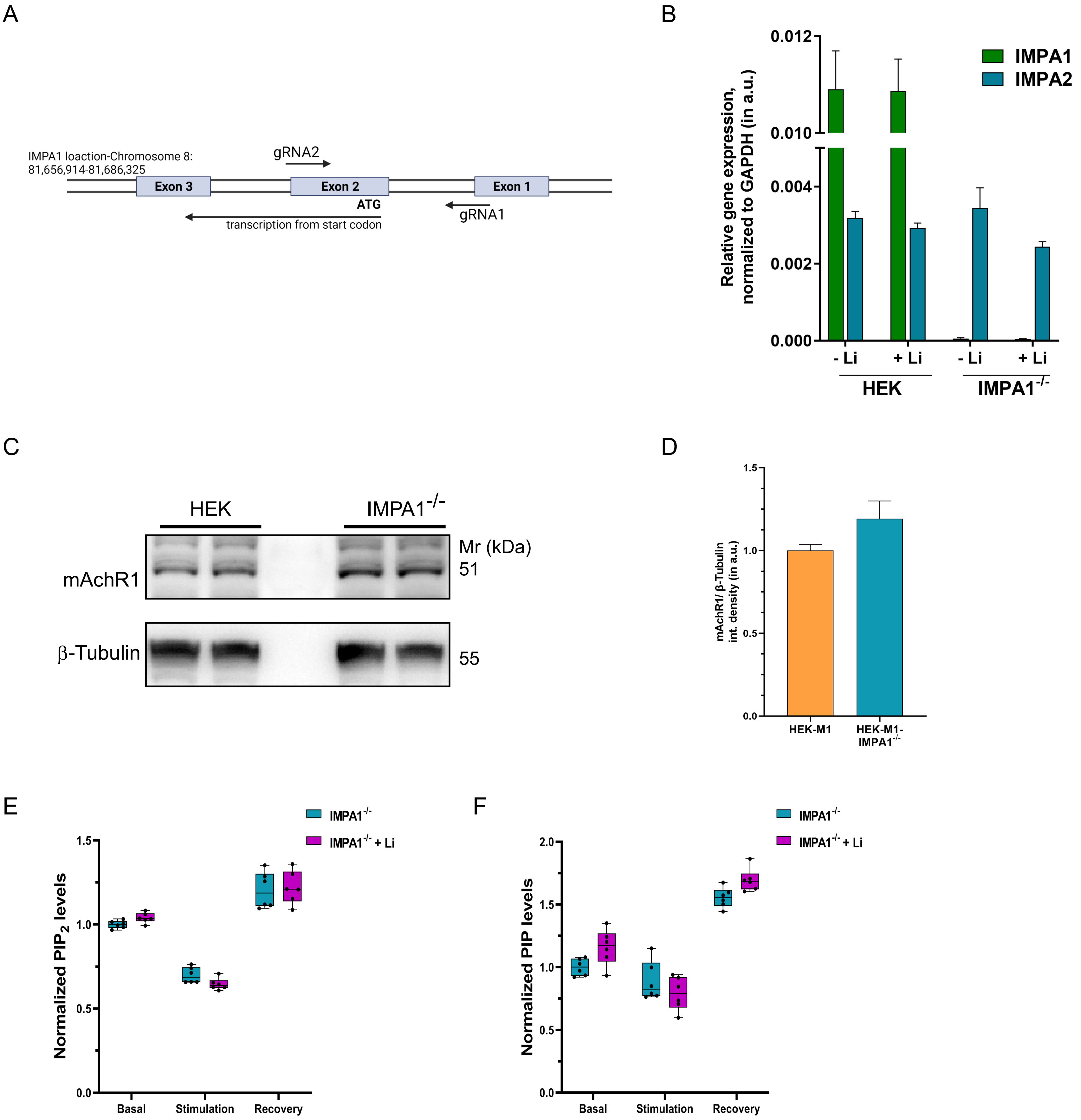


Saha Supp Figure 3

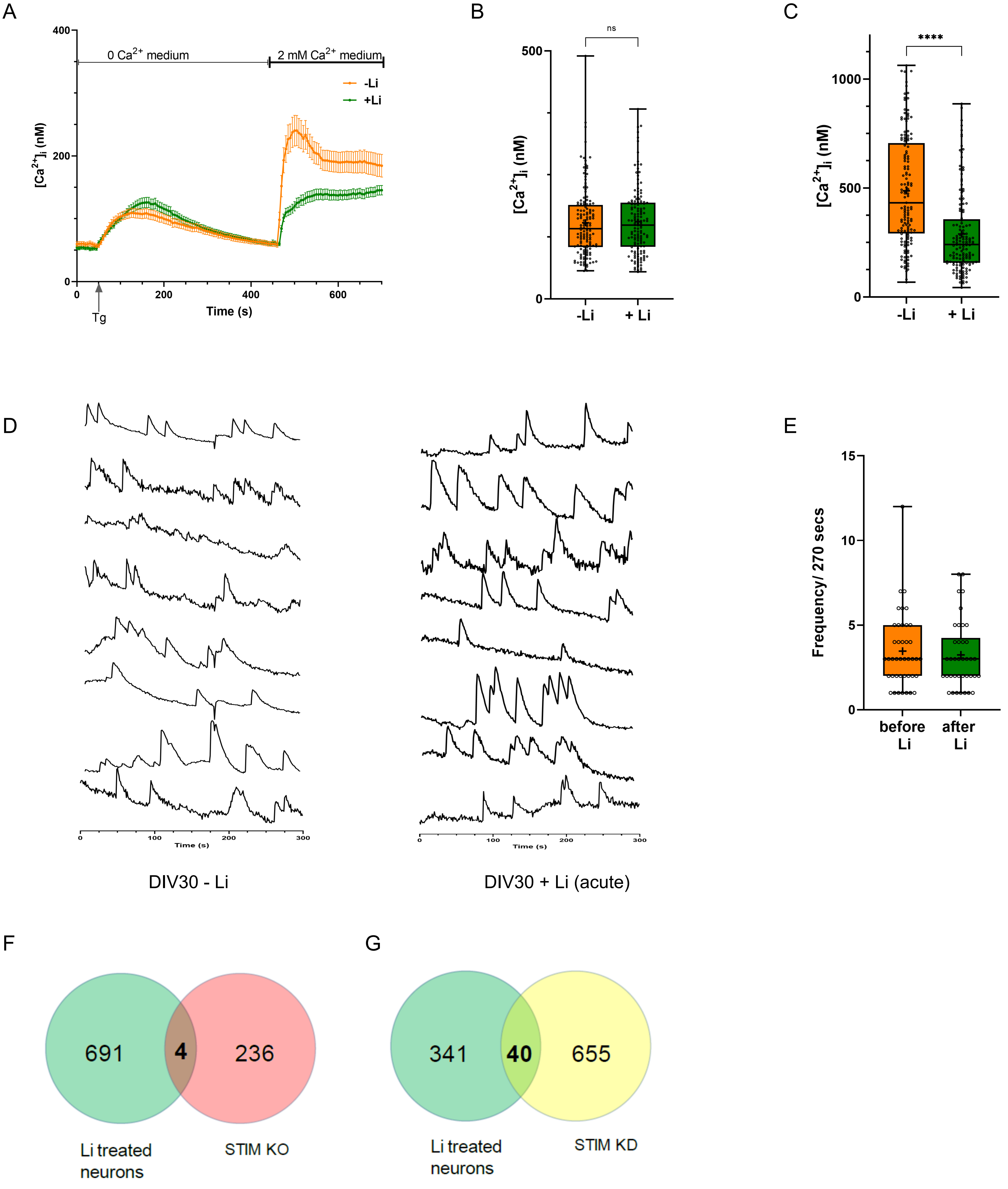


Saha Supp Figure 4

A

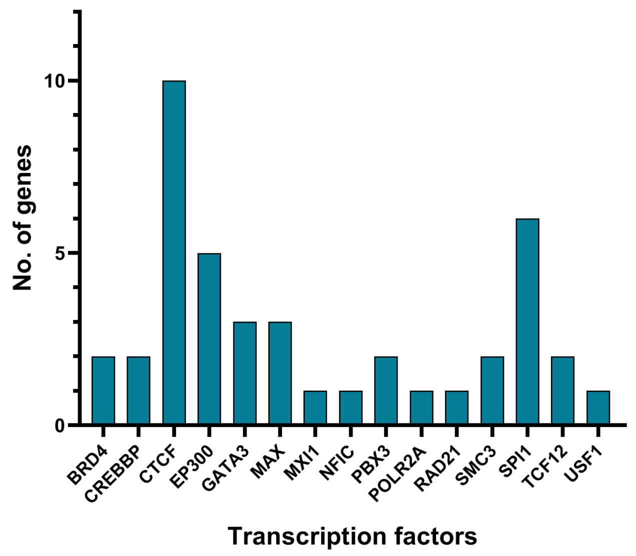


B

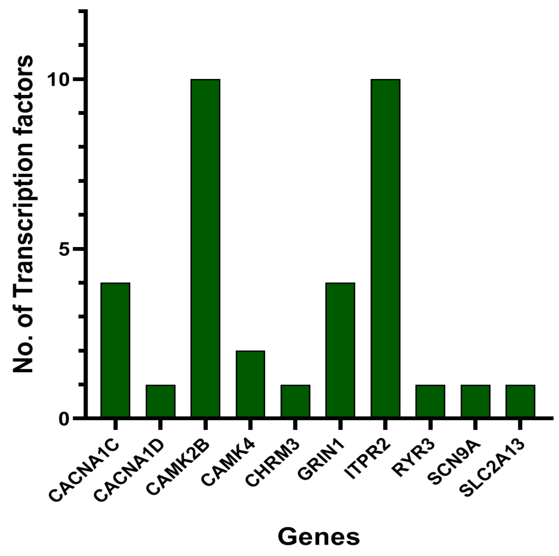


C

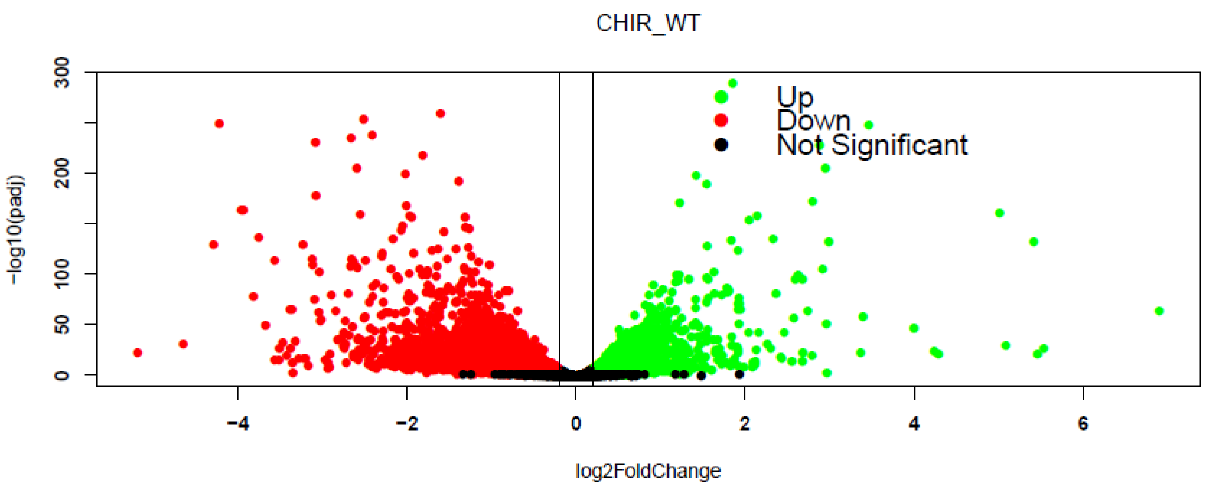


D

| Gene ID | Gene name | baseMean | log2 FoldChange | pvalue | padj |
|---|---|---|---|---|---|
| ENSG00000110092 | cyclinD1 | 1317.99 | 0.28 | 0.00 | 0.00 |
| ENSG00000078018 | MAP2 | 48889.94 | -0.14 | 0.00 | 0.01 |
| ENSG00000177606 | c-Jun | 4851.97 | 0.15 | 0.00 | 0.02 |
| ENSG00000141510 | P53 | 571.73 | 0.24 | 0.00 | 0.05 |
| ENSG00000080815 | Presenilin1 | 1901.77 | 0.15 | 0.01 | 0.09 |
| ENSG00000103197 | TSC2 | 2096.37 | -0.15 | 0.01 | 0.10 |
| ENSG00000070018 | LRP6 | 4202.20 | 0.14 | 0.01 | 0.11 |
| ENSG00000198561 | δ-catenin | 8187.44 | 0.09 | 0.02 | 0.13 |
| ENSG00000134982 | APC | 7104.11 | -0.10 | 0.04 | 0.20 |
| ENSG00000082701 | GSK3B | 0.00 | 0.08 | 0.04 | 0.22 |
| ENSG00000149294 | NCAM | 32954.36 | -0.06 | 0.08 | 0.30 |
| ENSG00000178403 | Neurogenin2 | 286.20 | 0.20 | 0.15 | 0.44 |
| ENSG00000162992 | NeuroD | 722.22 | 0.11 | 0.17 | 0.46 |
| ENSG00000142192 | APP | 37645.74 | 0.04 | 0.19 | 0.49 |
| ENSG00000197971 | Myelin basic protein | 386.66 | -0.12 | 0.23 | 0.55 |
| ENSG00000118513 | c-Myb | 132.05 | -0.16 | 0.33 | 0.64 |
| ENSG00000168036 | β-catenin | 11020.97 | -0.03 | 0.37 | 0.68 |
| ENSG00000171862 | PTEN | 4465.42 | -0.04 | 0.37 | 0.68 |
| ENSG00000118260 | CREB | 2453.46 | 0.05 | 0.40 | 0.71 |
| ENSG00000161202 | Dvl | 3265.50 | 0.03 | 0.42 | 0.72 |
| ENSG00000186868 | Tau | 27952.23 | -0.03 | 0.45 | 0.74 |
| ENSG00000103126 | Axin | 958.57 | 0.05 | 0.50 | 0.77 |
| ENSG00000170365 | SMAD1 | 738.14 | 0.05 | 0.51 | 0.78 |
| ENSG00000111361 | eIF2B | 1238.25 | 0.03 | 0.68 | 0.88 |
| ENSG00000092964 | CRMP2 | 31852.85 | -0.01 | 0.78 | 0.92 |
| ENSG00000074054 | CLASP1 | 6068.76 | -0.01 | 0.83 | 0.94 |
| ENSG00000131196 | NFAT | 68.44 | -0.04 | 0.87 | 0.96 |
| ENSG00000131711 | MAP1B | 98292.60 | 0.01 | 0.87 | 0.96 |
| ENSG00000162946 | DISC1 | 117.71 | -0.03 | 0.87 | 0.96 |
| ENSG00000164885 | CDK5 | 445.92 | -0.01 | 0.88 | 0.96 |
| ENSG00000130294 | Kinesin | 29314.79 | 0.00 | 0.94 | 0.98 |
| ENSG00000136997 | c-Myc | 437.65 | 0.00 | 0.98 | 1.00 |

E

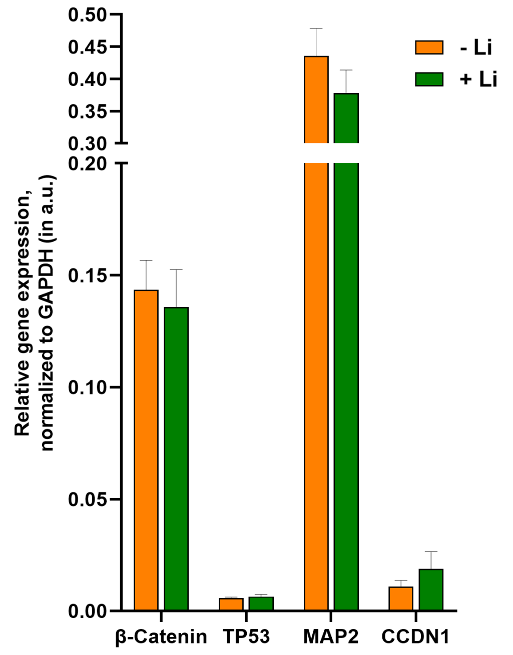


Saha Supp Figure 6

**Supplementary Figure 1- An in vivo model system to study the impact of Li on PLC induced $PIP_2$ turnover:**

(A) Nir2 downregulation due to shRNA. (B) Rate of regeneration of $PIP_2$ at the plasma membrane is shown for cells transfected with Nir2 shRNA, compared to cells transfected with scrambled shRNA. Mean ± SD is plotted from two experiments, each performed in replicates control cells (orange line, n =18), and for cells treated with Nir2 shRNA (blue line, n =21). (C, D) Relative abundance of the $PIP_2$ and PIP species shown, as detected by LC-MS (MRM method in the positive mode). [Statistical test: (A) Student's unpaired t test. * p value < 0.05; ** p value <0.01; *** p value < 0.001.]

**Supplementary Figure 2- Chronic Lithium treatment reduces PLC dependent intracellular $Ca^{2+}$ mobilization:**

(A-C) Expression studies were done to check whether the decrease in SOCE in the lithium treated HEK293T cells was due to downregulation of the genes involved in SOCE. No significant alteration occurred in terms of the expression of ITPR, STIM and ORAI in HEK293T cells due to lithium treatment.

**Supplementary Figure 3- IMPA1 is required for the effect of Li on $PIP_2$ resynthesis and agonist mediated $Ca^{2+}$ signalling:**

(A) The starting region of the CDS of *IMPA1* gene is targeted by guide RNAs (gRNAs) and deleted by spCas9; (B) *IMPA2* expression is not altered in these knock-out cells. *IMPA1* is the predominant gene in HEK293T cells that encodes for IMPA. (C, D) Western blot showing that both the reporter lines (HEK-M1 and HEK-IMPA1$^{-/-}$-M1) exhibit similar expression of the m1AchR and thereby should achieve similar PC stimulation due to same concentration of Cch. (E, F) Total $PIP_2$ and PIP levels using LCMS from whole cell lipid extract of untreated and lithium treated HEK IMPA1$^{-/-}$ cells (n=6 for both). [Statistical test: (E, F, H) Student's unpaired t test. * p value < 0.05; ** p value <0.01; *** p value < 0.001].

**Supplementary Figure 4- Lithium reduces excitability in human cortical neurons:**

(A-C) Intracellular store measured by Tg, is unaltered by lithium treatment in human cortical neurons. Quantification: untreated neurons (orange), n =145; neuronal cultures treated with

Li (green), n =145. (D) $[Ca^{2+}]_i$ traces from individual cells (30 DIV neurons) pre- and post-acute lithium treatment (1 mM Li for 5 mins) (soma; Y-axis shows ΔF and X-axis is time in seconds). Initially, the baseline was recorded for 300 secs- then 1 mM Li was added to the bath, incubated for 5 minutes and then recorded again for 300 secs. (E) The number of spikes/ 270 secs are counted from individual soma and plotted, each dot representing events from a single soma. Quantification- neurons before Li treatment (orange), n =41; neurons treated with Li for 5 mins (green), n =38. (F) The number of differentially expressed genes that are common between lithium treated neurons and in mouse Purkinje neurons where STIM1 is knocked out. (G) The number of differentially expressed genes that are common between lithium treated neurons and in neuronal progenitor cells where STIM1 is knocked down. [Statistical tests: (B, C) Student's unpaired t test. * p value < 0.05; ** p value <0.01; *** p value < 0.001; **** p value < 0.0001].

**Supplementary Figure 6- Lithium treatment induces transcriptional changes in pathways involved in neuronal transmission:**

(A) Transcription factors that regulate genes involved in calcium signalling were identified using hTFtarget database. The graph represents number of calcium signalling genes controlled by each transcription factor in the lithium treated neurons. (B) The graph represents number of transcription factors that control each gene involved in calcium signalling. (C) Volcano plot showing expression of genes altered due to CHIR99021 treatment ($p_{value}$ <0.05, $p_{adj\ value}$ <0.01). (D, E) Transcriptomic analyses were done to check whether Li treatment in mature neuronal affected genes involved in the Wnt signalling/ GSK-3β pathway. No significant alteration occurred in terms of the expression of the relevant genes due to lithium treatment.
